# CRMP/UNC-33 maintains neuronal microtubule arrays by promoting individual microtubule rescue

**DOI:** 10.1101/2024.05.31.596870

**Authors:** Xing Liang, Regina Agulto, Kelsie Eichel, Caitlin Ann Taylor, Victor Alexander Paat, Huichao Deng, Kassandra Ori-McKenney, Kang Shen

**Affiliations:** Department of Biology, Stanford University, Stanford, CA 94305, USA; Howard Hughes Medical Institute, Department of Biology, Stanford University, Stanford, United States; Department of Molecular and Cellular Biology, University of California, Davis, Davis, CA 95616, USA

**Keywords:** neuronal microtubule, microtubule associated proteins, CRMPs, microtubule dynamic instability, neuronal polarity

## Abstract

Microtubules (MTs) are intrinsically dynamic polymers. In neurons, staggered individual microtubules form stable, polarized acentrosomal MT arrays spanning the axon and dendrite to support long-distance intracellular transport. How the stability and polarity of these arrays are maintained when individual MTs remain highly dynamic is still an open question. Here we visualize MT arrays *in vivo* in *C. elegans* neurons with single microtubule resolution. We find that the CRMP family homolog, UNC-33, is essential for the stability and polarity of MT arrays in neurites. In *unc-33* mutants, MTs exhibit dramatically reduced rescue after catastrophe, develop gaps in coverage, and lose their polarity, leading to trafficking defects. UNC-33 is stably anchored on the cortical cytoskeleton and forms patch-like structures along the dendritic shaft. These discrete and stable UNC-33 patches concentrate free tubulins and correlate with MT rescue sites. *In vitro*, purified UNC-33 preferentially associates with MT tips and increases MT rescue frequency. Together, we propose that UNC-33 functions as a microtubule-associated protein (MAP) to promote individual MT rescue locally. Through this activity, UNC-33 prevents the loss of individual MTs, thereby maintaining the coverage and polarity of MT arrays throughout the lifetime of neurons.

## Introduction

Neurons are long-lived, highly polarized cells with complex morphologies. In axons and dendrites, long distance intracellular transport is mediated by the well-organized, parallel, staggered MT arrays ^1–3^. Neuronal MTs have several distinct features, including specialized polarity and stability. First, axonal and dendritic MTs have distinct polarities. In axons, MT plus ends are predominantly oriented in the distal direction, whereas in dendrites, the MT orientation is mixed, with a significant proportion of MTs adopting a minus-end-out configuration. The polarity of MT arrays is preserved throughout the lifespan of neurons^4–8^. In axons, MT polarity is maintained through dynein-mediated MT sorting and augmin-mediated MT nucleation using existing MTs as a template ^9^. However, less is known about how MT polarity is maintained in dendrites. Second, there exists a stable population of neuronal MTs that is both cold and drug resistant, properties that have been attributed to the binding of different microtubule-associated proteins (MAPs), supported by *in vitro* and *in vivo* evidence^10,11^. Although knock out mice of some MAPs show reduced MT density in axons or dendrites, progress in understanding the physiological function of the MAPs has been slow because of the potential redundant functions *in vivo* ^12^. Third, despite the overall stability of MT arrays, individual MTs exhibit dynamic growth and shrinkage in both axons and dendrites. Interestingly, there is electron microscopy evidence of stable and labile domains on individual MTs, possibly due to different tubulin posttranslational modifications or the association of different MAPs ^13–15^. For example, Tau is shown to preferentially bind MT labile domains and protect the MT dynamic ability in cultured rat neurons, which is contrary to its canonical function as MT stabilizer^14^. Understanding the interplay and relative contributions of stable and dynamic MT segments is crucial to unraveling the intricacies of MT array regulation within neurons. Defining MAPs that are essential for MT integrity *in vivo* will shed light on their functional significance in neuronal development, connectivity, and maintenance.

Collapsin response mediator proteins (CRMPs) were identified as MAPs and have been shown to play important roles in axon formation, guidance, synapse maturation and neuronal degeneration ^16–25^. These physiological functions of CRMPs in the nervous system are likely driven by regulating MT dynamics, however, the precise mechanisms are unknown ^23,26,27^. UNC-33, the lone CRMP homolog in *C. elegans*, is important for neuronal polarity in the PVD polymodal sensory neuron ^28^. A recent study showed that UNC-33/CRMP together with UNC-44/Ankyrin restricts MT sliding in PVD ^29^, which might contribute to the MT polarity phenotypes that manifest in the mutants ^28,29^. A separate line of research showed that CRMPs are some of the most stable proteins in the brain ^30^, so CRMPs might have a long-lasting function to maintain the MT organization during the lifespan of neuron. However, no direct evidence links the physiological function of CRMP family members to their activity in MT dynamic regulation.

In this study, we visualized the MT array in the *C. elegans* PVD sensory neuron with single MT resolution *in vivo*. We found that MTs formed a continuous, overlapping polarized array in the dendritic shaft. Within this array, individual MTs were highly dynamic, but consistently exhibited “rescue” events immediately following catastrophe. Furthermore, MT rescues in both the dendrite and axon were dependent on the activity and presence of UNC-33/CRMP. Intriguingly, UNC-33 localization along the neurite correlated with sites of rescue events. Loss of UNC-33 caused an increase in MT instability, shortening or complete loss of individual MTs, formation of gaps in the MT array and subsequently, disruption of MT polarity. Taken together, we found that UNC-33/CRMP maintains the integrity and polarity of the MT array by promoting individual MT rescues at specific sites *in vivo*, highlighting the essential role for this long-lived protein in neuronal organization and function.

## Results

### Dendritic microtubules are rescued at stereotyped locations

To understand the mechanisms that regulate the generation and maintenance of MT arrays in neurites *in vivo*, we used the sensory PVD neuron in *C. elegans* as a model system. After its birth, PVD sequentially extends an axon, followed by anterior and posterior primary dendrites. Higher order branches subsequently elaborate from the primary dendrites to form a regular menorah-like dendritic arbor (Fig. S1A) ^31,32^. We previously showed that the anterior primary dendrite contains exclusively minus-end-out MTs, and this polarized MT array is established by a developmentally active, mobile <u>d</u>endritic <u>g</u>rowth cone <u>MT</u> <u>o</u>rganizing <u>c</u>enter (dgMTOC), which localizes in the growth cone^33^. After the dendrite finishes its growth, the dgMTOC disappears, but MTs along the dendrite are maintained throughout the life of the neuron ^33^. To investigate how the MT array is organized and maintained in the absence of an apparent MTOC, we fused GFP to one of the α-tubulin isoforms, TBA-1, and expressed this construct in PVD neurons in a *tba-1* mutant background to reduce the free tubulin background signal. In the PVD cell body, GFP::TBA-1 exhibited a dynamic filamentous pattern, suggesting that TBA-1 was incorporated into polymerizing MTs (Fig. 1A). Similar to purified MTs, kymograph analysis of TBA-1 dynamics revealed distinct polymerization/growth and depolymerization/shrinkage events of GFP::TBA-1 labeled MTs *in vivo* (Fig. 1A’). Additionally, the non-uniform, stripe-like fluorescence pattern observed in the kymograph suggests that the incorporation of TBA-1 into MTs is not uniform, possibly due to the co-existence of other α-tubulins expressed in PVD neurons ^34^ (black and brown arrowheads in Fig. 1A’). In the mature primary dendrite, GFP::TBA-1 showed a continuous staining, consistent with complete coverage of *C. elegans* axons and dendrites by MTs ^33,35^. In contrast to the primary dendrite, only a fraction of the higher order dendritic branches contain MTs in PVD (Fig. 1B and 1B’), consistent with the results from Drosophila dendrites that some of the finest dendritic arbors lack MTs^36^.

**Fig. 1.**
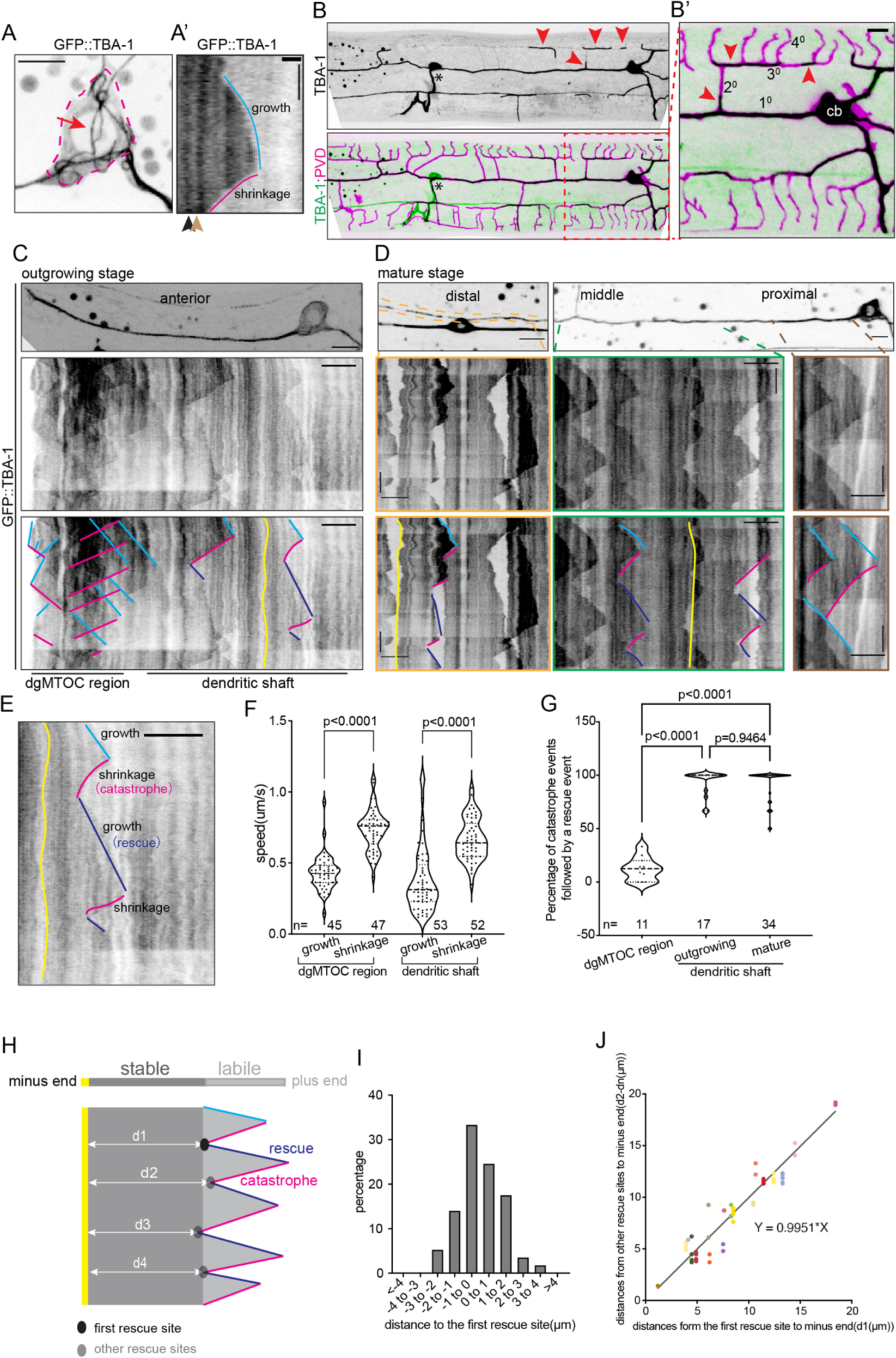
Dendritic microtubules are rescued at stereotyped locations. A) GFP::TBA-1 localization in PVD cell body(A) and kymograph of GFP::TBA-1 dynamic(A’). red arrow, microtubule filament labeled by GFP::TBA-1; black arrowhead, the bright stripe in kymograph; brown arrowhead, the dark stripe in kymograph. Blue line, MT polymerization; Magenta line, MT depolymerization; Red dashed line, outline of PVD cell body. B) GFP::TBA-1 localization in PVD anterior dendrite. Red arrowheads, microtubules in high order branches; black aster, HSN neuron. B’) Zoom in of the GFP::TBA-1 localization in the proximal region. C) GFP::TBA-1 localization (upper) and kymograph (lower) in an outgrowing anterior dendrite. D) GFP::TBA-1 localization (upper) and kymograph(lower) in mature primary dendrites. PVD distal dendrite is the process located in the orange dotted lines in the left panel. Orange boxes, distal region; green boxes, middle region, brown boxes, proximal region. E) Representative kymograph of MT dynamic behaviors in dendritic shaft. Lines in C-E: Light blue lines, growth events; magenta lines, shrinkage events; dark blue lines, rescue events; yellow lines, microtubule minus ends. Lines in lower panels of C) and D) are drown on the original kymograph of middle panels. F) Quantification of MT growth or shrinkage speed in MTOC region and dendritic shaft. P value is calculated by unpaired Student’s *t*-test. G) Quantification of the percentage of catastrophe events that are followed by a rescue event in different regions. P value is calculated by Brown-Forsythe and Welch ANOVA tests. H) Schematic of MT with stable and labile domains. Yellow region, MT minus end, dark gray region, MT stable domain, light gray region, MT labile domain, blue lines, MT growth or rescue events, magenta line, MT catastrophe events. d1-d4, distance between MT minus end to the different rescue sites. Black dot, the first rescue site; grey dots, other rescue sites. I) Distribution of distances between each other rescue sites and the first rescue site. n=57 rescue events from 18 animals. J) Quantification of the distance from each rescue site to MT minus end. Each color presents distances measured from same MT (n=18). The microtubule used for quantifying the distance between minus ends and rescue sites are the middle and distal microtubules with an identified minus end. Scale bar, 5μm for distance, 10s for time. See also Fig. S1.

To understand the process by which the MT array becomes organized and how the dynamic of individual MTs contribute to the formation of the MT array, we developed imaging protocols that allowed us to analyze both the collective MT organization and the dynamics of individual MTs with GFP::TBA-1. By taking advantage of the low number of MTs in *C. elegans* neurons^35^, we examined the MT dynamics in both developing and mature stages. During the outgrowth stage, continuous GFP::TBA-1 signal was found along the PVD primary dendrite, suggesting that a continuous MT array formed as the dendrite grew (Fig. 1C, upper). At this stage, TBA-1 dynamics visualized by kymograph analysis reveals two features of MT dynamics in dendrites. First, similar to their dynamics in the cell body, dendritic MTs showed both growth (Fig.1C, blue lines) and shrinkage (Fig.1C, magenta lines) behaviors. The transition from growth to shrinkage represents MT catastrophe (Fig. 1C, magenta lines) while the transition from shrinkage to growth represents rescue events (Fig.1C, dark blue lines). Second, the dynamic behaviors of MTs were different depending on their locations within the dendrite. Strikingly, MTs originating from the dgMTOC region only showed growth and shrinkage behavior with no observable rescues, while MTs in the rest of the dendrite (dendrite shaft) showed many rescue events despite having similar growth and shrinking speeds as those in the dgMTOC region (Fig. 1E to G). In summary, a continuous MT array forms from the dgMTOC during the outgrowth of dendrite, however, MTs within the dendritic shaft experience frequent rescue events along the lattice, unlike dgMTOC MTs in the growth cone region.

Next, we investigated how the MT array is maintained in the mature dendrite as the dgMTOC is a transient structure that only exists during dendrite outgrowth to establish the polarized MT array^33^. Maintaining MT coverage and uniform polarity for the life of the neuron requires the longevity of MT filaments at the level of individual MTs. This could be achieved by stabilizing MTs and completely freezing MT dynamics, or by ensuring consistent MT rescue events after catastrophe. To distinguish between these possibilities, we analyzed TBA-1 dynamics over time in mature dendrites. Given the differences in MT dynamics within the dgMTOC region and dendritic shaft during development, we divided the mature primary dendrite into three segments: proximal, middle, and distal segments (Fig. S1A and Fig.1D). Using GFP::TBA-1 kymographs, we observed robust MT dynamics in all three segments of the mature dendrite. We identified the growth (blue) and catastrophe (magenta) phases for individual MTs and quantified the percentage of catastrophe events which were followed by a rescue event within the mature dendrite. These results show that mature dendritic MTs display consistent (∼100%) rescue events, similar to the dendritic shaft MTs during outgrowth (Fig. 1G). This indicates that MTs undergo frequent growth and shrinking cycles with consistent rescues events in mature dendrites, which underlies the longevity of MTs.

Although certain MAPs and factors like tubulin concentration and GTP-bound tubulin islands (GTP islands) can promote rescues *in vitro*, little is known about the features and mechanisms of MT rescue in axons and dendrites ^37–40^. In PVD dendrites, we often observe multiple rescue events from the same MT in one imaging experiment (Fig. 1H). The low MT number (1-2MTs) in the middle and distal dendritic shaft made it possible to identify a single MT and determine its dynamic plus end and stable minus ends based on GFP::TBA-1 fluorescence (yellow lines in Fig. 1C to E). The identification of minus ends was further confirmed by the localization of the MT minus end protein, PTRN-1^41–43^ (Fig. S1B). With the identification of both minus ends and rescue sites, we plotted the distance from each rescue site to the first rescue site, and the distance from each rescue site to the minus end, respectively. For MTs with 2 or more rescue events, we found that most of the rescue events of the same MT occurred within 1 micron along individual MTs (Fig. 1H and I), regardless of the length of MT (Fig. 1J). The stereotyped rescue sites suggest that each dendritic MT has a relatively stable domain and a labile domain (Fig. 1H), which has also been observed in axonal MTs ^44,35^.

To further validate the stable and labile domains, we expressed the rigor Kinesin-1 mutant UNC-116(G237A) which binds preferentially to stable MTs ^45^. UNC-116(G237A) localized to neurites and the cell body of PVD and marked a fraction of somatic MTs labeled by GFP::TBA-1 (Fig. S1C). Time lapse experiments show that UNC-116(G237A) is largely bound to the MT stable domain but not the labile domain, which further supports the notion of stable and labile domains for individual MTs (Fig. S1D). Together, these data indicate that the dendritic MTs are dynamic but experience reliable rescue events that occur at stereotyped locations along the lattice (Fig. 1H). It is conceivable that consistent rescue events are essential for maintaining MT polarity and number, because once MTs are lost in mature dendrites, the dgMTOC is no longer present to either generate new MTs or provide templated polarity.

To examine the MT stability over a longer time period, we tagged TBA-1 with a photoconvertible fluorescent protein, pcStar, which switches from green to red upon exposure to ultraviolet (UV) light. After illuminating a small region of the primary dendrite with a UV laser for 1ms, the converted region became visible in the red channel, and the green unconverted signal was reduced in the same area (Fig. S1E). The same dendrite was imaged one hour after conversion, and we found that about 60% of the red signal in the converted region remained (Fig. S1F), indicating that the stable portion of MTs lasted for at least one hour.

### CRMP/UNC-33 is required for maintaining the continuous MT array

While it is widely accepted that MAPs play critical roles in regulating MT dynamics, the precise molecules essential for MT array stability in neurites are unknown. Taking advantage of our ability to directly visualize MTs in neurites, we performed a candidate genetic approach to identify MAPs that affect formation of the continuous MT array in PVD. We found that *unc-33* mutants showed the most dramatic MT distribution phenotype. *unc-33* encodes the sole *C. elegans* homolog of the vertebrate collapsing response mediator proteins (CRMPs)^46^. In both *unc-33(mn407)* null and missense *unc-33(e204)* loss of function mutants, we observed a fully penetrant phenotype where gaps developed in the MT array (∼2-3 gaps/100µm), with an average gap length of 19.43±11.71μm in the null mutant compared to almost no gaps in wild-type animal (Fig. 2A-C). UNC-33 was recently shown to form a complex with UNC-119/UNC119 and UNC-44/Ankyrin. This tripartite complex anchors MTs and prevents them from sliding ^29^. Therefore, we further tested *unc-44* and *unc-119* mutants and found that both mutants showed similar gap phenotypes as *unc-33* mutants, suggesting that this anchoring complex is important for the continuous MT array formation, maintenance, or both (Fig. 2A-C).

**Fig. 2.**
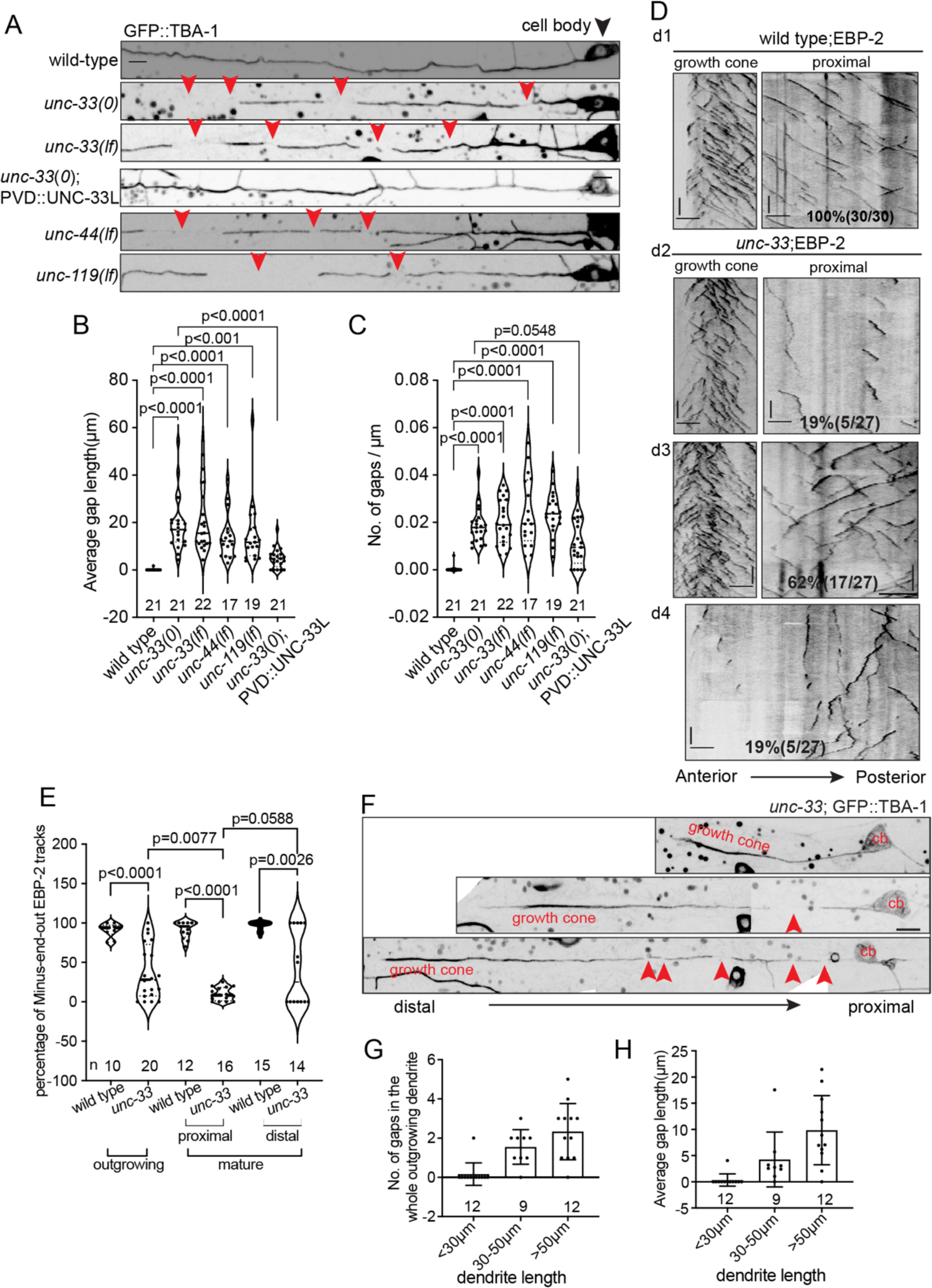
UNC-33/CRMP is required for maintaining the continuous MT array. A) GFP::TBA-1 localization in the proximal region of mature dendrites in wild-type, *unc-33(0)*,*unc-33*(*lf*), *unc-33(0);* PVD::UNC-33L, *unc-44(lf)* and *unc-119(lf)* mutants. Red arrowheads, MT gaps in primary dendrites. Black arrowhead, PVD cell body. B) Quantification of gap number. (C) Quantification of average gap length. D) Representative kymograph of EBP-2 dynamic in outgrowing wild-type(d1) and *unc-33* mutants PVD dendrites(d2-d4). E) Percentage of minus-end-out EBP-2 tracks in wild-type and *unc-33* mutants in outgrowing and mature dendrites. F) GFP::TBA-1 localization in outgrowing dendrites of *unc-33* mutants. Red arrowheads, gaps in MT arrays of outgrowing dendrites; cb, cell body. G) Quantification of gap number in outgrowing dendrites with different length. H) Quantification of average gap length in outgrowing dendrites with different length. P values are calculated by Brown-Forsythe and Welch ANOVA tests. N number is labeled below each column. Scale bar, 5μm for distance, 10s for time. See also Fig. S2 to Fig. S4.

Next, we aimed to understand how the gaps arise in the MT array. In PVD, a single dgMTOC generates new MTs and establishes MT polarity for the dendritic shaft during development ^33^(d1 in Fig. 2D). Given that previous studies have suggested a role for UNC-33 in MT polarity in mature PVD dendrites ^28^, we therefore examined if UNC-33 affected MT array formation by regulating dgMTOC localization or organization using endogenously tagged MT plus end tracking protein EBP-2::GFP. During PVD dendrite outgrowth, ∼81% of *unc-33* mutants showed a normal, active dgMTOC in the growth cone region based on EBP-2 dynamics (d2 and d3 in Fig. 2D). The remaining mutants (∼19%) of the mutants did not have a dgMTOC and therefore displayed abnormal dendrite MT polarity (d4 in Fig. 2D). However, within *unc-33* mutants with normal dgMTOC in the growth cone, the majority (∼76%) showed a mixed MT polarity in the proximal dendrite, which is different from the consistent minus end out MT polarity in the proximal dendrite of wild type animals (d3 in Fig. 2D), indicating that the lack of dgMTOC might not be the direct cause of the MT polarity defects in *unc-33* mutants. We next considered if the perturbations in MT polarity was a result of a maintenance defect that arises after the MT array is established by the dgMTOC. This is supported by the observation that MT polarity defects observed in proximal dendrites of *unc-33* mutants are further exacerbated in mature dendrites (Fig. 2E). In the distal dendrite, a population of *unc-33* mutants showed reversed MT polarity while the remaining mutants displayed normal MT polarity, likely reflecting the partial penetrance of dgMTOC defects in this mutant (Fig. 2E and Fig. S2A).

To directly test if UNC-33 is required to maintain MT polarity after development, we examined the consequence of UNC-33 degradation after the completion of primary dendrite outgrowth using AID-mediated protein degradation system. We fused the degron sequence to the endogenous UNC-33, and then introduced the expression of TIR1 using a ubiquitous *eft-3* promoter. To bypass the early effect of UNC-33 in MT organization, we treated the strains with 4mM auxin for 18h after the primary dendrite finished growing. After treatment, the worms with both the degron-tagged UNC-33 and TIR1 expression showed a dramatic increase in plus-end-out MTs, suggesting that *unc-33* is required to maintain MT polarity (Fig. S2B and C). We also noticed that a small portion of worms showed polarity defects before auxin treatment, which is likely caused by the protein tagging and leaky degradation (Fig. S2B and C). Together, these data support the notion that the MT polarity defects in *unc-33* mutants are unlikely to be caused by a lack of a dgMTOC but instead result from a MT maintenance defect.

To directly test if the MT gaps observed in *unc-33* mutants arise from an inability to maintain the MT array, we examined the MT array directly using GFP::TBA-1 at different stages of dendrite outgrowth in *unc-33* mutants. Consistent with the polarity defect, MT gaps increase in both the number and length as the dendrite grows longer (Fig. 2F-H). This data further supports the notion that UNC-33 plays an essential role in the maintenance of the MT array.

Next, to test if the function of UNC-33 is conserved across different types of neurons, we examined the MT organization in the motoneuron DA9. In wild-type animals, GFP::TBA-1 showed a continuous localization in both the axon and dendrite in DA9. This is consistent with previous evidence using serial electron microscopy reconstruction^35^. In *unc-33* mutants, DA9 showed numerous gaps in both axons and dendrites, suggesting that UNC-33 functions in diverse types of neurons (Fig. S2D-F). Taken together, we found that the microtubule-associated protein, UNC-33, is required to maintain the intact MT array after it is established from the dgMTOC.

To understand how the polarity and gap defects arise, we recorded MT dynamics by performing time lapse imaging in the proximal dendrite, where MT gaps form first. Kymograph analysis showed that the gap formation was tightly correlated with MT dynamics, including MT depolymerization and sliding. However, MT sliding events were bidirectional which can both create and cover gaps, while MT depolymerization consistently causes gaps (Fig. S3A). While recording MT dynamics in *unc-33* mutants, we also observed an increase in plus end out MTs growing into the anterior dendrite from the cell body when there was a gap in the MT array (middle and right panels in Fig. S3A). This provides one potential mechanism to explain how MT polarity is disrupted in *unc-33* mutants since these cell body-derived MTs are of opposite polarity to the existing dendritic minus end out MTs. Consistent with this idea, the MT polarity in the proximal posterior dendrite in *unc-33* mutants was normal despite the presence of MT gaps in the posterior dendrite, since the endogenous MTs in the posterior dendrites are plus end out (Fig.S3B and C). In fact, Kinesin-2 has been proposed to stop the growth of plus-end-out MTs into dendrites by generating pulling force against the existing minus end out MTs and, therefore, maintain the minus-end-out MT polarity in *Drosophila* dendrites ^47^. Whether the increase in cell body-derived plus end out MTs is a direct consequence of gap formation or other MT dynamics defects in *unc-33* mutant needs to be further tested.

### UNC-33 promotes microtubule rescue

To understand how UNC-33 regulates MT dynamics, we compared MT dynamics *in vivo* with single MT resolution in wild type and *unc-33* mutants using GFP::TBA-1. We found that *unc-33* mutants showed normal polymerization and depolymerization speeds, length and dynamicity at both plus ends and minus ends (Fig. S3D-M). However, *unc-33* mutant MTs showed dramatically reduced rescue frequency (Fig. 3A-C) while the catastrophe frequency was normal (Fig. S3N). In wild type animals, the vast majority of catastrophe events were followed by a rescue event in the dendritic shaft (Fig. 1C-H and Fig. 3A). However, in *unc-33* mutants, a significant portion of catastrophe events showed no subsequent rescues, causing complete depolymerization of the MT (Fig. 3B, D and E), likely leading to reduced MT number and increased MT gaps. Consistent with the previously reported anchoring function of UNC-33 ^29^, we also observed sliding of individual MTs in *unc-33* mutants, suggesting that UNC-33 has multiple roles in regulating MT dynamics (Fig. 3B and E). Furthermore, *unc-44* mutants showed the same increased frequency of MT loss and sliding, further suggesting that the UNC-33/UNC-119/UNC-44 complex functions together to promote MT rescue and anchoring (Fig. 3E).

**Fig. 3.**
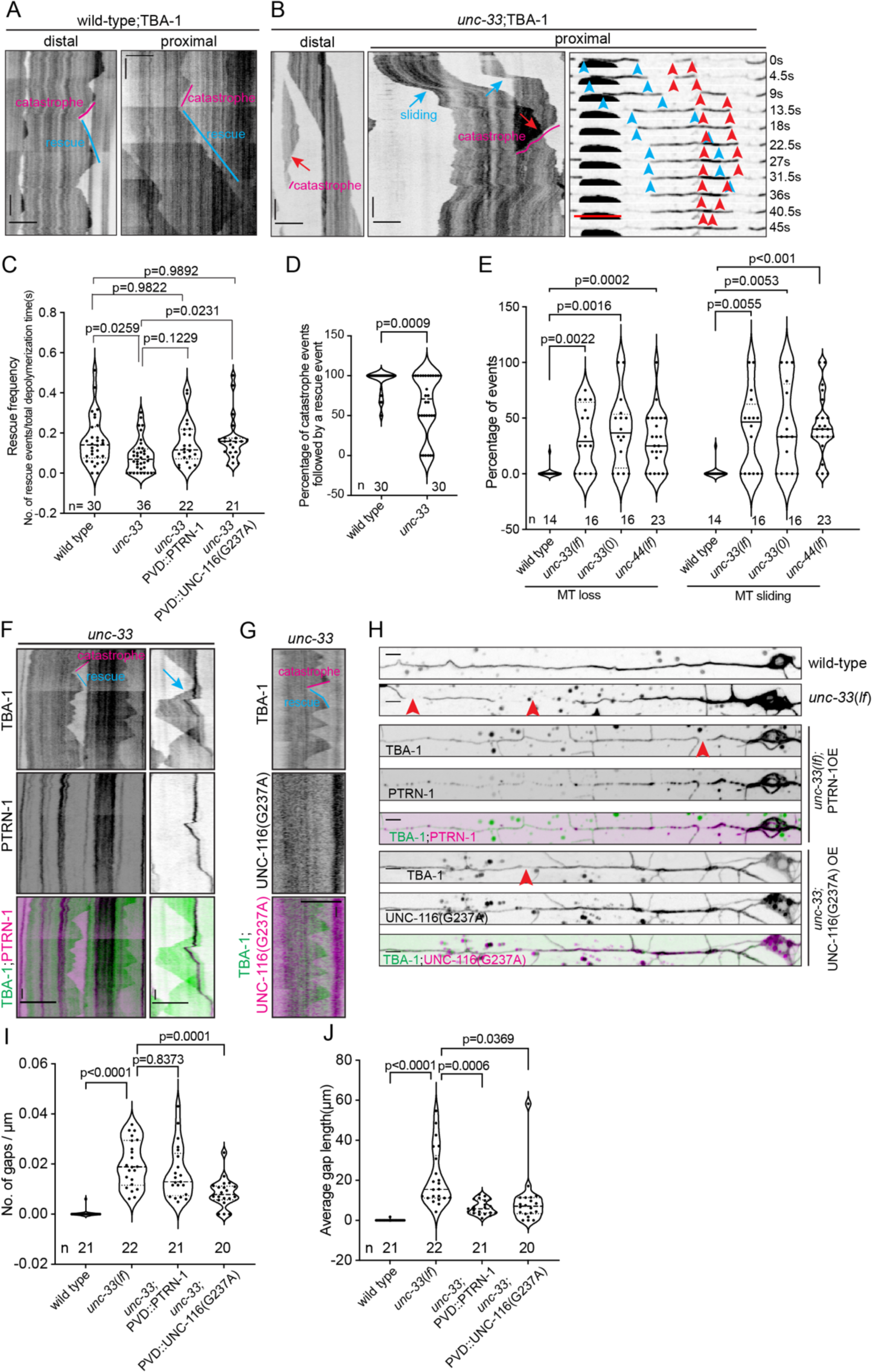
UNC-33 promotes MT rescue. A) Representative kymograph of GFP::TBA-1 in wild-type animal. Magenta lines, catastrophe events; blue lines, rescue events. B) Representative kymograph of GFP::TBA-1 in *unc-33* mutants. Magenta lines, catastrophe events; red arrows/arrowheads, MT loss, blue arrows/arrowheads, MT sliding. C) Quantification of MT rescue frequency in animals of different genotypes. D) Quantification of the percentage of catastrophe events that are followed by a rescue event in wild type and *unc-33* mutants. The quantification data for wild type are from the same animals in Fig. 1G. E) Quantification of MT loss and MT sliding in animals of different genotypes. F) Representative kymograph of GFP::TBA-1 and PTRN-1::tdtomato dynamics in *unc-33* mutants. G) Representative kymograph of GFP::TBA-1 and UNC-116(G237A)::mCherry dynamics in *unc-33* mutants.. For F) and G), Magenta lines, catastrophe events; blue lines, rescue events; blue arrow, MT sliding. H) GFP::TBA-1 localization in proximal dendrites of animals with different genotypes. Red arrowheads, gaps in MT array. I) Quantification of gap number. J) Quantification of average gap length. Scale bar, 5μm for distance, 10s for time. P values are analyzed by Brown-Forsythe and Welch ANOVA tests. See also Fig. S2 to Fig. S4.

To determine whether the MT gaps are the result of MT loss or sliding, we used known molecular manipulations to stabilize MTs in *unc-33* mutants. PTRN-1 is a MT minus end binding protein which stabilizes MT minus ends ^41–43^. PTRN-1 can also bind along the MT and stabilize MTs when overexpressed^41^. In *unc-33* mutants, overexpressed PTRN-1 formed puncta along the dendrite (Fig. 3F). During MT sliding, PTRN-1 puncta moved synchronously with the MT, indicating that overexpressed PTRN-1 binds along MTs (Fig. 3F, blue arrow). Interestingly, overexpressing PTRN-1 in *unc-33* mutants significantly increased MT rescue frequency, and rescue events often occurred at PTRN-1 puncta (Fig.3C and F). In contrast, the frequency of MT sliding events in the PTRN-1 overexpressing worms was unchanged compared to *unc-33* mutants (Fig. S3O). Coincidently, PTRN-1 overexpression also dramatically reduced the gap length in *unc-33* mutants, but did not significantly affect the number of gaps significantly (Fig. 3H-J). Together, these data indicate that MT instability is the main cause of the MT gap formation in *unc-33* mutants. To further confirm the role of MT rescue in MT array gap formation, we examined whether overexpression of the “rigor” form of Kinesin-1 rescues the gap phenotype in *unc-33* mutants, as growing evidence has shown that the binding of Kinesin-1 plays a role in expansion and stabilization of the MT lattice ^48–50^. We showed that the rigor mutant Kinesin-1 specifically binds to the stable domains of MT filaments in wild-type animals (Fig. S1D). When the rigor form of *C. elegans* Kinesin-1, UNC-116(G237A), was overexpressed in *unc-33* mutants, we found that the MT rescue frequency was completely rescued, indicating that UNC-116(G237A) stabilizes individual MTs *in vivo* (Fig. 3C and 3G). Interestingly, overexpression of UNC-116(G237A) also reduced the number and length of gaps, further supporting the notion that reduced MT rescue events in the *unc-33* mutant are responsible for the gap phenotype (Fig. 3H-3J). Taken together, we show that UNC-33 maintains the intact MT array in dendrites by promoting individual MT rescue.

To further confirm UNC-33 stabilizes MTs, we performed a photoconversion experiment in which we converted the pcStar::TBA-1 from green to red in the *unc-33* mutant background. While the red signal remained stable the in wild type animal, in *unc-33* mutants the converted red signal was dramatically reduced one hour after conversion (Fig. S4A and B), suggesting that MT stability is reduced in *unc-33* mutant.

Since we have shown that UNC-33 is required for maintaining MT polarity, we next investigated whether MT rescue activity contributes to polarity maintenance. We induced polarity maintenance defects by degrading UNC-33 using auxin induced degradation and used PTRN-1 overexpression to stabilize MTs and asked if the polarity phenotype could be rescued. Indeed, PTRN-1 overexpression partially rescued the MT polarity defect in auxin treated animals (Fig. S2C). It is worth noting that PTRN-1 puncta along the dendrite protected short MT seeds and occasionally acted as MT nucleation sites, which caused additional polarity defects by promoting MT growth in random directions (Fig. S4C). This effect potentially underestimates PTRN-1’s ability to rescue the MT polarity phenotypes in *unc-33* mutants. A previous study proposed that MT sliding could cause polarity deficits^29^, however, our PTRN-1 overexpression did not rescue the MT sliding phenotype (Fig. S3O). This is an expected result because PTRN-1 is not connected to UNC-44 like UNC-33. Based on the partial rescue of the polarity phenotype by PTRN-1 overexpression, it is plausible that both the MT rescue activity and the MT anchoring activity of UNC-33 contribute to polarity maintenance.

### UNC-33 localizes to a patch-like structure to promote microtubule rescue in the dendrite

The CRMP family of proteins are conserved MAPs, which bind to both tubulins and MTs *in vitro*^18,24,25,51,52^. To understand how UNC-33 promotes MT rescue and maintains MT stable domains *in vivo*, we first studied the expression and subcellular localization of endogenous UNC-33. The *unc-33* gene contains three alternatively spliced isoforms. Expression of cDNA encoding the long isoform rescues the MT gap phenotype in *unc-33* mutants (Fig. 2A-C), suggesting that the long isoform is functional. We found that insertion of a tag at either the N or C terminus of the *unc-33* endogenous gene resulted in uncoordinated worms similar to the *unc-33* mutants, suggesting that these tags disrupted UNC-33 function (^28,29^ and data not shown). To circumvent this problem, we inserted the GFP internally in the long isoform of UNC-33, which resulted in animals with normal locomotion (Fig. S5A). Using this functional endogenous UNC-33::GFP strain, we found UNC-33 was expressed broadly in many tissues, including PVD (Fig. S5B), which is consistent with its cell-autonomous function^28^. UNC-33::GFP appears to localize to patch-like structures along neurites (Fig. S5B). However, its broad expression in many neurons made it difficult to study UNC-33’s subcellular localization in PVD. Therefore, we knocked in a 3xGFP11 fragment in the same genomic locus and combined it with GFP1-10 specifically expressed in PVD to achieve single cell endogenous labeling of UNC-33 (Fig. S5A). The PVD specific labeling strain showed that UNC-33 localizes to patch-like structures along PVD dendrites, both in primary and higher order dendrites (Fig. 4A). Consistent with its function in both the axon and dendrite, we also observed that UNC-33 forms similar patch-like structures in the axon of PVD and FLP, another highly branched sensory neuron in *C. elegans* ^53^ (Fig. S5C).

**Fig. 4.**
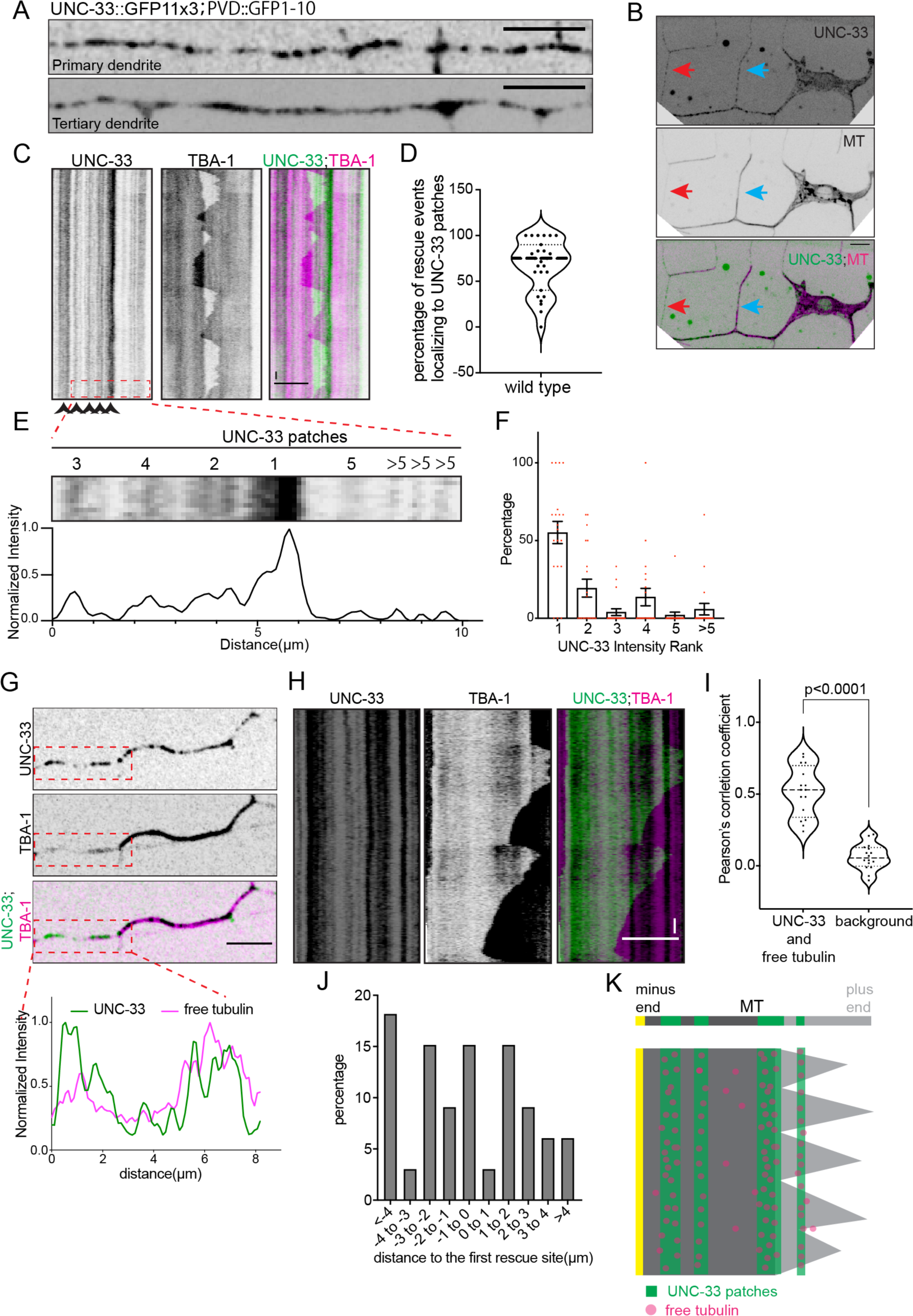
UNC-33 localizes to a patch-like structure to promote MT rescue in dendrites. A)UNC-33 localization in PVD primary and tertiary dendrites. B) UNC-33 and TBA-1 localization in PVD. Red arrows, branches with UNC-33 but not MT; blue arrows, branches with both UNC-33 and MT. C) Representative kymograph of UNC-33:(int)GFP and mCherry::TBA-1 dynamic in PVD tertiary dendrites. D)Quantification of the percentage of rescue events that localize to UNC-33 patches in tertiary dendrites (n=27 individual MTs). E) An example of UNC-33 patches ranking by the maximum intensity. F) Distribution of percentage of the rescue events happening in the UNC-33 patches of different intensity. n=20 rescue events from 12 animals. G) UNC-33 and TBA-1 localization in PVD tertiary dendrite (upper), and Normalized intensity of UNC-33 and TBA-1 intensity in the MT free region of tertiary dendrite(lower). Red box, MT free region of tertiary dendrite. H) Representative kymograph of UNC-33 and TBA-1 dynamic in PVD tertiary dendrite which shows the free tubulin localization in a MT free region. I) Quantification of the colocalization between UNC-33 and free tubulin signal in MT free regions in tertiary dendrites. n=16 MT free regions. P value is analyzed by Welch’s t test. J) Distribution of distances between each other rescue site and the first rescue site in *unc-33* mutants. n=33 rescue events from 16 MTs. K) Schematic model of how UNC-33 promotes MT rescue at stereotyped locations: UNC-33 localizes to patch-like structures along dendrites which are stable, and these stable structures promote MT rescue by concentrating free tubulins, thus proving stereotyped rescue sites for MTs. Scale bar, 5μm for distance, 10s for time. See also Fig. S5 and S6.

To understand how UNC-33 promotes MT rescue, we visualized UNC-33 and TBA-1 simultaneously and found that UNC-33 localized to all the branches, while MTs only entered a subset of branches, suggesting that the localization pattern of UNC-33 is not dependent on MTs (Fig. 4B). We further recorded MT dynamics while imaging UNC-33 localization at tertiary branches where there are only one or two MTs. The kymographs showed that UNC-33 patches are largely stationary, unlike the dynamic growth and shrinkage of MTs. In areas where only one MT exists, the UNC-33 patch did not disappear upon MT depolymerization (Fig. 4C). This supports the hypothesis that the localization pattern of UNC-33 is not dependent on MTs, which is also consistent with a recent report that UNC-33 is anchored by the UNC-119 and UNC-44 complex to the plasma membrane^29^. Importantly, we identified each MT rescue event and asked whether the rescue events coincided with an UNC-33 patch. This analysis showed that the majority of MT rescue events happened at UNC-33 positive regions (Fig. 4C and D), indicating that UNC-33 promotes MT rescue locally. While the density of UNC-33 patches is high, we noticed a correlation between the intensity of individual UNC-33 patches and MT rescues (Fig.4C). To further quantify this correlation, we ranked the UNC-33 patches near MT rescue sites by their intensity in each kymograph and calculated the rescue percentage at each patch for the same MT (Fig. 4E). This analysis showed that the majority of rescue events happen at the brightest UNC-33 patch, suggesting that MT rescue activity is dependent on UNC-33 concentrations (Fig. 4F).

Next, we asked how UNC-33 promotes MT rescue in a concentration-dependent manner. An important structural study showed that CRMP2’s binding affinity to the GTP form of tubulin-dimers is much higher than its affinity to MTs, and that the CRMP2-tubulin dimer binding promotes MT assembly^51^. It is therefore plausible that the UNC-33 patches in PVD dendrite could recruit and increase the local concentration of free tubulin dimers as a mechanism to increase rescue frequency. To directly test this hypothesis, we took advantage of the fact that MTs are not present along the entire length of tertiary dendrites at all times (Fig. 1B and B’).We performed time-lapse experiments to unambiguously identify MT-free regions because the free tubulin signals are much lower compared to that of a single MT (red box in Fig.4G). In the MT-free region, we observed that the tubulin signal is not evenly distributed. Instead, higher tubulin concentration correlates with UNC-33 patches (Fig. 4G-I). This observation is consistent with the possibility that UNC-33 locally concentrates tubulins and promotes MT rescue by generating a local tubulin pool. Through this mechanism, the stationary UNC-33 patches define stereotyped locations of rescue events and demarcate the borders of the stable MT domain. Consistent with this hypothesis, we observed MT rescue from random sites in *unc-33* mutants (Fig. 4J and K).

To directly test whether UNC-33 concentrated free tubulins, we artificially targeted the short isoform UNC-33 to the mitochondria outer membrane by fusing a mitochondria-targeting sequence to the unc-33 cDNA (Mito-UNC-33S). The short isoform lacks the N terminal binding domain required for the UNC-44 and UNC-119 complex, but contains the same C terminal MT binding domain and the tubulin binding domain as the long isoform. Mitochondria form a tubular network in the PVD cell body and discrete puncta along the primary dendrite^54^. With the overexpression of mito-UNC-33S, we observed concentrated tubulin signal that showed strong colocalization with mitochondria (Fig. S6A), indicating that UNC-33S was sufficient to create local tubulin pools. Further, we introduced the mito-UNC-33S transgene into the *unc-33* mutant and observed similar concentrated tubulin signals in both the cell body and dendrite (Fig. S6B) Dynamic imaging in the MT-free dendritic region in *unc-33* mutants showed that no obvious MT dynamic behavior occurred in the mito-UNC-33S region, demonstrating that mito-UNC-33S concentrated tubulins rather than binding and recruiting MTs (Fig. S6B). To examine whether the mito-UNC-33S could promote MT rescue in an *unc-33* mutant background, we performed dynamic timelapse imaging in a dendritic region with both MT and mito-UNC-33S signal, and we observed occasional MT rescue events at the mito-UNC-33S site (Fig. S6C). We did not observe repeated rescue events at the same sites, possibly due to the high level of MT sliding (Fig. S6C). Taken together, we propose that UNC-33 patches promote MT rescue by creating highly concentrated tubulin pools in dendrites.

To further test the causality between the UNC-33 patches and MT rescue, we sought to disrupt UNC-33’s subcellular localization. UNC-44/Ankyrin is likely required for UNC-33’s pattern because it forms a complex with UNC-33 to inhibit MT sliding in the dendrite^29^. First, we examined the subcellular localization of the long isoform of UNC-44 with a knock in UNC-44::FLPon-mScarlet in combination with flippase-mediated recombination to achieve single cell endogenous labeling of UNC-44. This long isoform of UNC-44 showed dim fluorescence signal along the dendrite in addition to being enriched at the axon initial segment^55^. Within dendrites, UNC-44 showed a patch-like pattern, partially colocalizing with UNC-33::GFP (Fig. S7A). Further, in *unc-44* mutants, UNC-33 largely lost its patch-like appearance and became diffuse throughout the dendrite and cell body with an increased intensity in the cell body (Fig. 5A-C). In *unc-44* mutants, the MT rescue events were similarly reduced compared to *unc-33* mutants (Fig. 3E). When we observed rescue events of a single MT in *unc-44* mutants, the events occurred at random sites along the dendrite, but not at stereotyped locations (Fig. 5D, E and Fig. S7B). These results are consistent with the notion that the UNC-33 patches locally promote MT rescue and demarcate a stable domain for individual MTs.

**Fig. 5.**
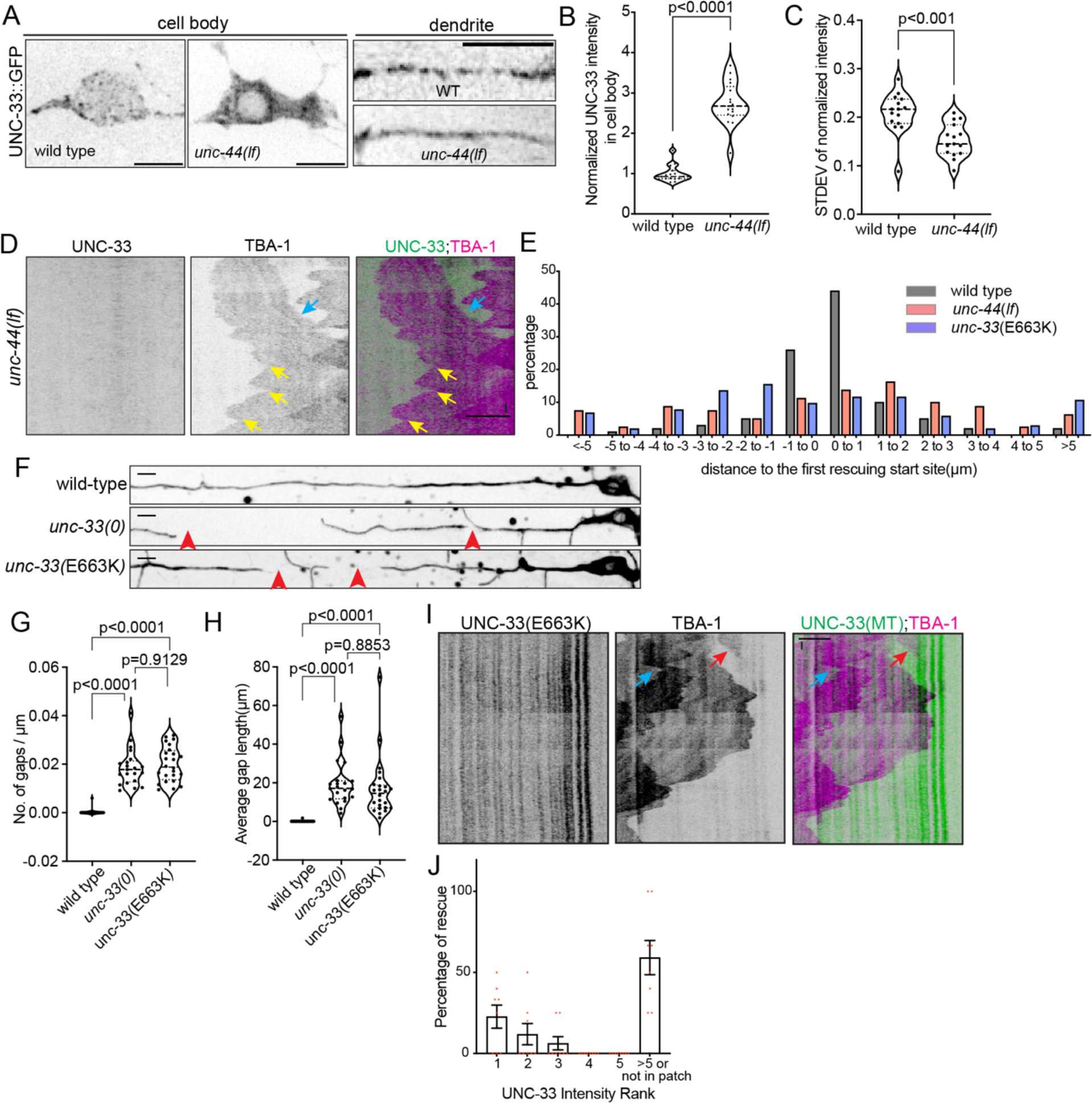
Both patch-like localization and MT assembly activity are required for UNC-33 to promote the stereotyped MT rescue. A) UNC-33 localization in cell body and dendrites in wild-type and *unc-44(lf)* mutants. B) Quantification of UNC-33 intensity in the cell body of wild type and *unc-44* mutants. n=15 for wild type, n=17 for *unc-44* mutants. P value is calculated by unpaired Student’s *t*-test with Welch’s correction. C) Quantification of the STDEV of normalized intensity of UNC-33 in wild-type and *unc-44(lf)* mutants. n=15 for wild-type and *unc-44* mutants. P value is calculated by unpaired t test with Welch’s correction. D) Kymograph of UNC-33:(int)GFP and mCherry::TBA-1 dynamic in PVD tertiary dendrites in *unc-44* mutants. Blue arrows, MT sliding, yellow arrows, rescue sites of the same MT. E) Distribution of distances between each other rescue sites and the first rescue site in wild type, *unc-44* mutants and *unc-33(E663K)* mutants. n=100 rescue events from 23 MTs in wild type animals, n=80 rescue events from 23 MTs in *unc-44* mutants, n=103 rescue events from 42 MTs in *unc-33*(E663K) mutants. F) GFP::TBA-1 localization in primary dendrites of wild-type, *unc-33*(*0*) and *unc-33*(E663K) mutants. Red arrowheads, gaps in MT array. G) Quantification of gap number in wild-type*(n=21)*, *unc-33*(*0*)(n=21), *unc-33*(E663K)(n=24) animals. H) Quantifications of average gap length in wild-type*(n=21)*, *unc-33*(*0*)(n=21), *unc-33*(E663K) (n=24) animals. For G) and H), P values are calculated by Brown-Forsythe and Welch ANOVA test. The quantification data for wild type and *unc-33*(0) mutants are the same as Fig. 2B and C. I) Kymograph of UNC-33(E663K):(int)GFP and mCherry::TBA-1 dynamic in PVD tertiary dendrites. Red arrows, MT loss; blue arrows, MT sliding. J) Distribution of percentage of the rescue events happening in the UNC-33(E663K) patches of different intensity. n=8 rescue events. Scale bar, 5μm for distance, 10s for time. See also Fig. S7.

To understand how UNC-33 promote MT rescue, we next disrupted its MT binding with a mutation in its MT binding domain. The UNC-33 homolog CRMP has been shown to promote MT polymerization through its C-terminal MT assembly domain, and the E663K point mutation in the conserved domain of *unc-33* showed MT polarity defects in neurites^28,56^. The same point mutant caused a similar MT gap phenotype as the *unc-33* null mutant, suggesting that the “MT assembly domain” of UNC-33 is important for the MT rescue function (Fig. 5F-H). To further understand how this mutation affects UNC-33 protein level, localization, or activity, we engineered the same point mutation in the 3xGFP11 endogenous UNC-33 background and combined it with expression of GFP1-10 specifically in PVD. We found that the mutation did not change the patch-like localization of UNC-33; however, the mutated UNC-33 failed to induce local MT rescue (Fig. 5E, I, J and Fig. S7C). Taken together, these results indicate that UNC-33 localizes to patch-like structures in an UNC-44 dependent manner. UNC-33 patches promote local MT rescue and delineate the stable and labile domain of individual MTs. UNC-33’s MT rescuing activity is essential for the formation and continuous MT array in neurites.

### UNC-33 promotes microtubule rescue *in vitro*

Having identified a role of UNC-33 in rescuing MTs *in vivo*, we next turned to *in vitro* experiments to directly examine whether UNC-33 affects parameters of MT dynamic instability, including polymerization, catastrophe, and rescue as the mammalian CRMPs have been shown to promote MT assembly *in vitro* ^17,56,57^. We recombinantly expressed and purified full-length, N-terminally sfGFP-tagged *C. elegans* UNC-33. Since the UNC-33 short isoform contains the conserved MT assembly and tubulin binding domains, we chose to express and purify the short form to test its interaction with MTs (Fig. 6A). We first measured the mass distribution of the purified sfGFP-UNC-33 by mass photometry and found that purified sfGFP-UNC-33 can exist as a monomer, but can also form a tetrameric complex (Fig. 6B), which is consistent with prior work showing that mammalian CRMP2 forms tetramers ^51^. Using total internal reflection fluorescence microscopy (TIRF-M), we imaged the binding and behavior of sfGFP-UNC-33 on taxol-stabilized MTs and found that while UNC-33 bound along the MT lattice, it preferentially associated and accumulated at MT ends (Fig. 6C). We then analyzed the behavior of sfGFP-UNC-33 on dynamic MTs. In this assay, we affix stable GMPCPP MT seeds to the coverslip, then image as free tubulin in solution polymerizes off the seed in either the absence or presence of sfGFP-UNC-33. Again, we observed sfGFP-UNC-33 accumulate at the plus-ends of growing MTs (Fig. 6D). We observed a modest increase in plus-end growth rate in the presence versus the absence of sfGFP-UNC-33 (Fig. 6E). Although we did not observe any significant difference in the catastrophe frequency in the absence versus presence of sfGFP-UNC-33, we did observe a significant 42% increase in rescue frequency in the presence of sfGFP-UNC-33 (Fig. 6E-G). Under our experimental conditions, we also noted that sfGFP-UNC-33 promoted active rescues within the GDP lattice following catastrophe events, whereas in its absence, MTs frequently depolymerized to the GMPCPP seed (Fig. 6D). In summary, our TIRF-M imaging data suggests that UNC-33 is sufficient to promote MT rescue *in vitro*.

**Fig. 6.**
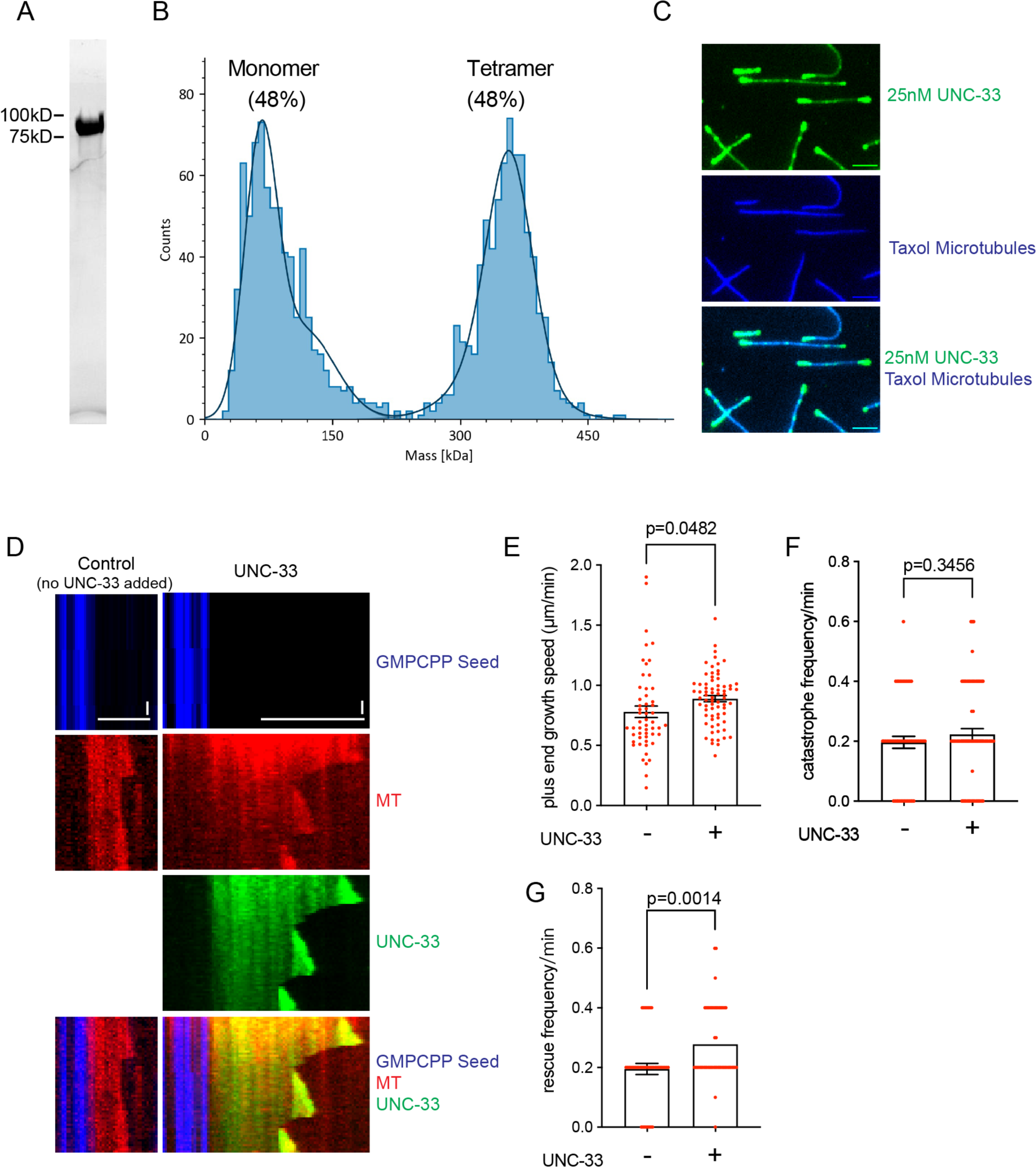
UNC-33 promotes microtubule rescue *in vitro*. A) SDS-PAGE gel of sfGFP-UNC-33 short isoform purified from BL21 cells. B) Mass photometry of sfGFP-UNC-33 (expected mass: kDa). Fits of mean mass values for monomer (observed mass: 77 kDa) and tetramer (observed mass: 355 kDa) species and relative fraction of particles with each indicated mass are shown. C) Representative TIRF-M images showing 25 nM of UNC-33 (green) binding to taxol-stabilized-microtubules (blue). Scale bars = 2 μm. D) Kymographs of microtubule dynamics showing the polymerization of tubulin (red, 15 μM) off of GMPCPP seeds (blue) in the absence or presence of sfGFP-UNC-33 (green). Scale bars: 5 μm for distance, 30s for time. E) Quantification of microtubule plus end growth speed in the absence (n=54) or presence (n=71) of UNC-33. F) Quantification of the catastrophe frequency speed in the absence (n=54) or presence (n=70) UNC-33. G) Quantification of microtubule rescue frequency speed in the absence (n=43) or presence (n=54) of UNC-33. All graphs display all data points with means and s.e.m. All p-values were calculated using a Welch’s t-test.

### CRMP/UNC-33 is excluded from the growth cone region

Since MTs nucleated from the dgMTOC showed much fewer rescue events when compared with MTs in the dendritic shaft (Fig. 1G), we asked whether this is caused by a region-specific localization of UNC-33 during development. To probe this question, we examined the subcellular localization of UNC-33 in PVD during the outgrowth of its primary dendrite. We first combined the UNC-33::3xGFP11 knock in worm with GFP1-10 driven by a promoter which is active in the outgrowing PVD. However, this approach did not yield sufficiently bright GFP for our microscopy experiment, likely due to the low expression level early in development. We next used the endogenously GFP::UNC-33 knock-in strain. As shown above, UNC-33::GFP knock-in is broadly expressed in many cells and tissues in *C. elegans* (Fig. S5B), which made it difficult to isolate the signal from the PVD growth cone. To eliminate the UNC-33 signal from other cells, we photobleached all UNC-33 signals in an area where the PVD dendrite was about to grow into. Since the PVD dendritic growth is stereotyped between animals and the only growing structure in the bleached area is the PVD dendrite, we can reliably visualize UNC-33::GFP signal together with the dendritic growth cone. Under this condition, we found that UNC-33 was excluded from the growth cone area but becomes present in the more proximal dendrite. This ∼15-20 µm segment from the dendritic tip matches the size of the dgMTOC region (Fig. 7A and B). To further examine UNC-33’s subcellular localization in PVD, we overexpressed mNeonGreen tagged UNC-33 driven by cell type-specific *unc-86* promoter. Overexpressed UNC-33 showed similar localization as the endogenous labeled UNC-33 (Fig. 7C and D), demonstrating the exclusion of UNC-33 from the growth cone. Therefore, UNC-33 is absent from the dgMTOC region where MTs are highly dynamic with no rescue events. This is critical for MT polarity establishment during development as the plus end out MTs nucleated by the dgMTOC need to disappear to ensure a minus end out MT polarity during the movement of the dgMTOC ^33^.

**Fig. 7.**
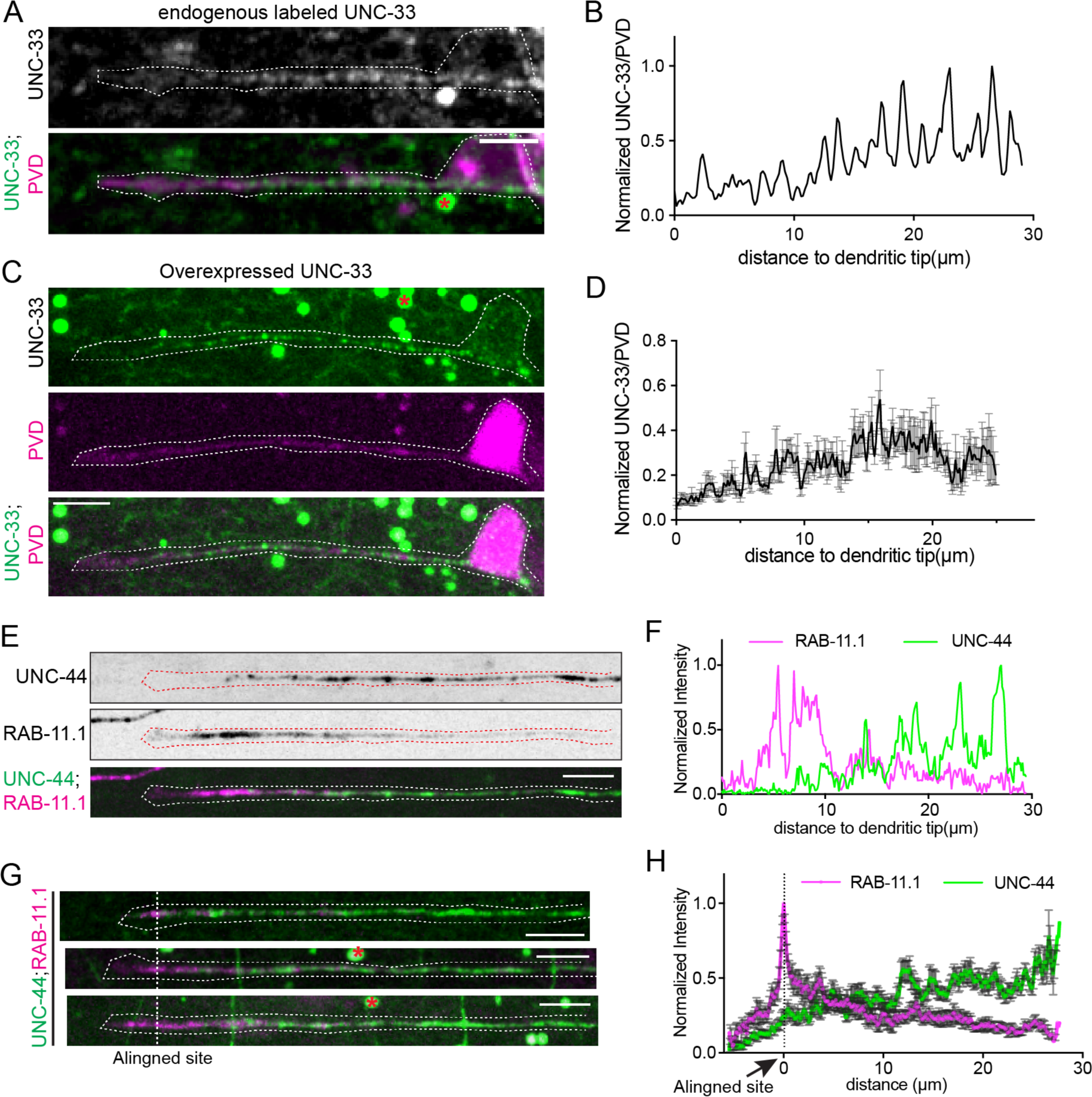
UNC-33/CRMP is excluded from the growth cone during development. A) Endogenous UNC-33:(int)GFP localization in PVD during development. White dashed lines, outlines of PVD morphology. B) Line distribution of normalized UNC-33 intensity along outgrowing primary dendrite. C) Overexpressed UNC-33::(int)mNeonGreen localization in PVD during development. (C) Line distribution of normalized UNC-33 intensity along outgrowing primary. n=7 animals. E) Endogenous UNC-44::GFP and PVD specific mCherry::RAB-11.1 localization in PVD primary dendrite during development. Red dashed lines, outlines of outgrowing PVD dendrites. F) Normalized intensity of UNC-44(green) and RAB-11.1(magenta) along primary dendrite during PVD development. G) Schematic of alignment of RAB-11.1 in the growth cone region. H) Normalized intensity of UNC-44(green) and RAB-11.1(magenta) along primary dendrite during PVD development. n=10 animas. Scale bar, 5μm. Red asters, auto fluorescent signals.

To further confirm this result and explore the localization between the dgMTOC and the anchoring complex, we examined the subcellular localization between UNC-44 and RAB-11.1, which labels endosomes and is localized to the dgMTOC ^33^. Interestingly, in developing dendrites, we observed complementary localization patterns between these two proteins. UNC-44 is largely absent from the most distal part of the PVD dendrite, where RAB-11.1 labeled endosomes reside. Trailing the dgMTOC area, UNC-44 showed patch-like structures along the dendrite shaft, similar to the localization of UNC-33 (Fig. 7E and F). To quantify the complementary localization between UNC-44 and RAB-11.1, we aligned the images from multiple worms to the brightest RAB-11.1 locus in the growth cone and plotted the intensity distribution of these two proteins, and we observed a strong UNC-44 signal where the RAB-11.1 signal was low (Fig. 7G and H). These results suggest that the UNC-33/UNC-44 complex continues to trail the mobile dgMTOC in the outgrowing dendrite, thus protecting the MTs after they are released from the dgMTOC. Taken together, our data suggest that UNC-33/UNC-119/UNC-44 specifically localizes to the developing and mature dendritic shaft and plays essential roles in the maintenance of the integrity, dynamic, and polarity of MT array in neurites.

### Continuous microtubule arrays are essential for long-distance trafficking in dendrites

To understand the consequence of a disconnected MT array with disrupted polarity, we examined the intracellular transport with several vesicle cargoes as MTs are intracellular highways for kinesin and dynein-mediated transport. First, we examined the synaptic vesicle marker RAB-3. Consistent with previous reports^28^, we found reduced RAB-3 localization in axons and ectopic localization of RAB-3 in dendrites in *unc-33* mutants, likely due to the increased plus end out MTs in the anterior dendrite (Fig. 8A and B). Moreover, we also noticed that the mislocalized RAB-3 in dendrites formed clusters along dendrites (Fig. 8A), which might be correlated with MT gaps. To directly examine the relationship between ectopic cargoes and MT gaps, we next examined a cargo that is normally localized to dendrites, RAB-11.1, which also showed abnormal distribution patterns along the PVD dendrites and tended to accumulate into clusters in *unc-33* mutants (Fig. 8C). For this experiment, we simultaneously imaged RAB-11.1 together with TBA-1 dynamics in PVD dendrites in wild-type animals and *unc-33* mutants. In wild-type animals, RAB-11.1 showed an even distribution along the anterior dendrite, and RAB-11.1 vesicles move seamlessly along the continuous MT array (Fig. 8C and D). In *unc-33* mutants, RAB-11.1 vesicles accumulated at the edges of MT gaps (Fig. 8C, right panel). Remarkably, RAB-11.1 vesicles start to move immediately after a growing MT bridges the gap (Fig. 8E). These data show that the continuous MT array is absolutely required for processive, long-distance transport. It also implies that molecular motors can be fully engaged at the end of MTs.

**Fig. 8.**
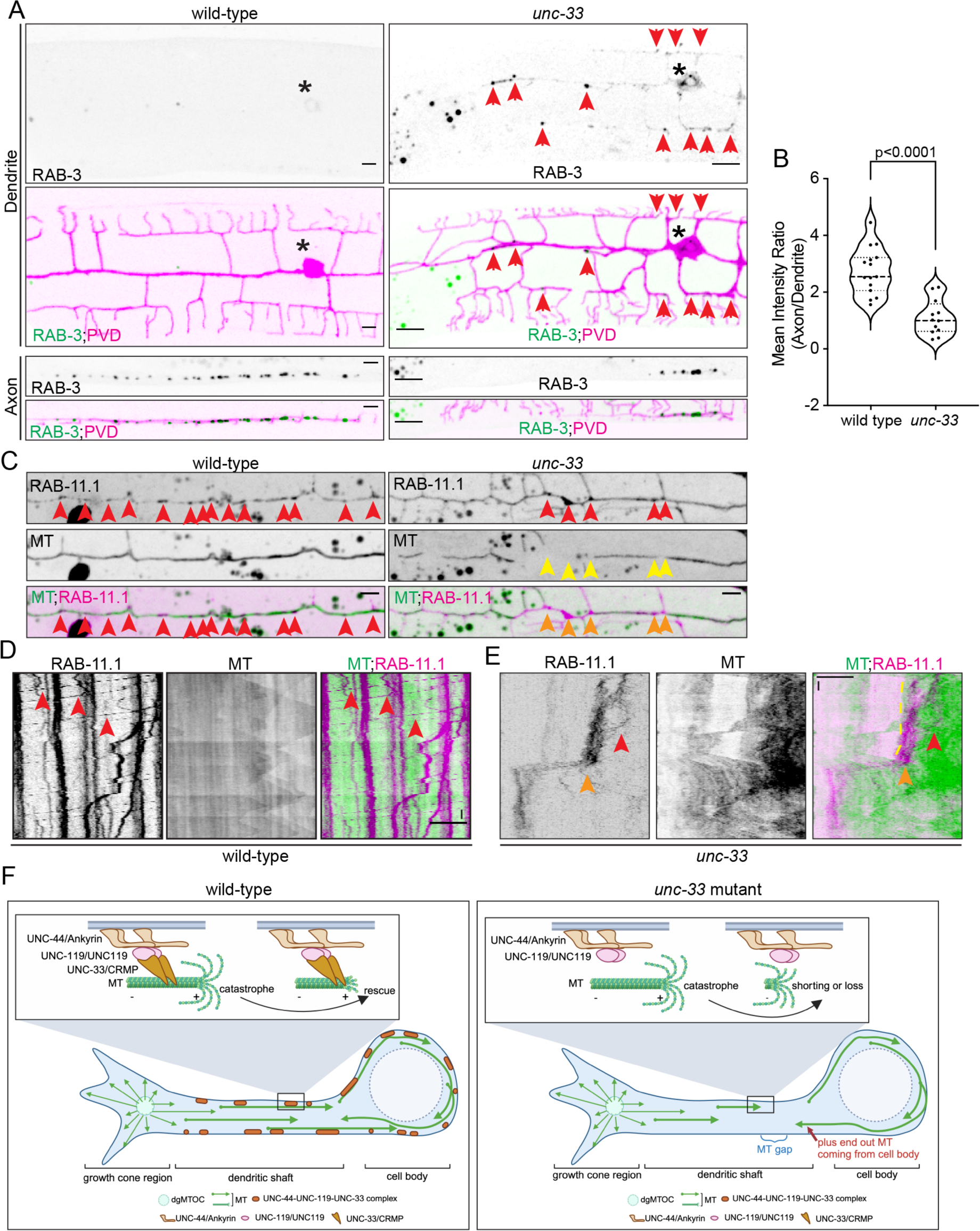
Continuous microtubule arrays are essential for long distance trafficking in dendrites. A) Endogenous RAB-3 localization in dendrites and axons of wild-type animals and *unc-33* mutants. Asterisk, cell body. Red arrows, RAB-3 positive vesicles that accumulated in dendrites in *unc-33* mutants. B) Quantification of polarized RAB-3 localization in wild-type (n=15) and *unc-33* mutants (n=12). ****p<0.0001, Welch’s t test. C) Localization of RAB-11.1 positive vesicles in dendrites of wild-type animals (left panel) and *unc-33* mutants (right panel). Red arrowheads, RBA-11.1 positive vesicles; yellow arrowheads, MT gaps; orange arrowheads, RAB-11.1 positive vesicles that accumulated at the MT gap sites. D-E) Kymograph of mCherry::RAB-11.1 and GFP::TBA-1 in primary dendrites in wild-type animals and *unc-33* mutants. Red arrowheads, continuous moving RAB-11.1 vesicles in primary dendrites. Orange arrowheads, accumulated RAB-11.1 positive vesicles at the MT gap sites. Yellow dashed line, the edge of the MT gap. F) Schematic model of the working mechanism of UNC-33. In wild-type animals, the continuous, polarized MT array established by dgMTOC is maintained by UNC-33/CRMP. UNC-33, which is anchored by UNC-44-UNC-119 complex, localizes to patch-like structures along dendritic shaft and these patches promote MT rescue locally. The stereotyped rescue events create the stable and labile domains of MTs and protect the MT coverage and polarity. Scale bar, 5μm for distance, 10s for time.

## Discussion

The unique MT organization in axons and dendrites is essential for efficient MT-based transport ^1,58^. Our previous study showed that in postmitotic neurons, the dgMTOC is responsible for the MT nucleation and polarity establishment during dendrite morphogenesis. This dgMTOC is active only during neurite outgrowth, but becomes inactive after neurite maturation^33^. We show here that the continuous, polarized MT array established by the dgMTOC is maintained by one of the conserved MAPs, UNC-33/CRMP. UNC-33 localizes to patch-like structures along the dendritic shaft, and these patches promote MT rescue locally by generating a local free tubulin pool. The stereotyped rescue events create the stable and labile domains of MTs and protect the MT coverage and polarity (Fig. 8F). These results not only uncovered key functions for UNC-33, but also unveiled a general framework for MT organization during the development and maintenance of dendrites.

First, our results shed light on how UNC-33 promotes MT rescue, maintaining a continuous MT array. Different from the canonical MAPs, the localization of UNC-33 is independent of MTs *in vivo*. Our mitochondria targeting experiment supports that UNC-33 patches recruit and increase the local concentration of tubulin *in vivo*. The last helix of CRMP2 has been shown to interact with GTP-bound tubulin dimers *in vitro*^51^, indicating that UNC-33 patches facilitate the formation of a tubulin pool enriched with GTP-bound tubulins. The GTP-bound tubulin pool might directly supply tubulin dimers for MT rescue. Alternatively, the local GTP-tubulin pool might favor the formation of GTP-rich domains along MTs, known as “GTP islands”, in nearby MTs. These GTP-tubulin islands in the middle of MTs slow down MT depolymerization and promote MT rescue^37,59^. *O*ur *in vitro* experiments further demonstrate that UNC-33 is sufficient to promote MT rescue. Since tubulin and UNC-33 are the only proteins in these reconstituted experiments, it provides evidence that UNC-33’s effects on MT rescue are direct. We note that UNC-33 does not prevent catastrophes neither *in vivo* nor *in vitro.* MT dynamicity is essential for many cellular processes^60^. UNC-33 ensures a continuous MT array without suppressing MT dynamics, providing both stability and dynamicity which is a unique feature for UNC-33 when compared with other MAPs^61–63^. In neurites, UNC-33 is anchored by the UNC-44/UNC-119 complex at the membrane where it determines the sites for MT rescue and forms the boundaries of the stable and labile domains for individual MTs.

Second, different dendritic regions show distinct MT dynamics. The dgMTOC region exhibits highly dynamic MTs with numerous polymerization and depolymerization events but few rescue events. In contrast, in the dendritic shaft, MTs are far less dynamic and exhibit consistent rescue events (Fig.1). The extreme dynamicity in the dgMTOC region is critical to maintain the minus-end-out MT polarity in dendrites during development^33^. However, after releasing from the dgMTOC, MTs in the dendritic shaft need to be protected to form a stable array. UNC-33 and UNC-44 are excluded from the dgMTOC region and only form patch-like structures along the dendritic shaft. This ensures that only minus-end-out MTs behind the dgMTOC will persist, hence protecting the MT coverage and maintaining the unidirectional polarity of MTs in the PVD anterior dendrite. Both UNC-119/UNC119 and UNC-44/Ankyrin have been shown to act as structural components to promote axon maturation by inhibiting membrane dynamics within the growth cone^64,65^. Therefore, the mechanisms of growth cone membrane dynamics might be intrinsically linked to MT dynamics through the localization of UNC-33 and UNC-44.

Third, MT stability is essential for the maintenance of MT polarity in dendrites. Because the dgMTOC is only present during the outgrowth phase, it can no longer provide a polarity template in mature dendrites. Interestingly, individual MTs exhibit reliable rescue events in the mature dendrite, which prevents complete depolymerization of MTs. This is essential for maintaining both the coverage of the MT array and the orientation of MTs in dendrites. In *unc-33* mutants, individual MTs undergo complete depolymerization and the loss of these MTs creates gaps in the MT array. It appears that with the gaps in the proximal dendrites, MTs from the cell body grow into dendrites and generate a plus-end-out MT population, which leads to MT polarity defects in the proximal dendrite. It is plausible that the unusually high MT stability in neurons is to prevent complete depolymerization events and subsequent loss of both mass and polarity.

Taken together, using PVD as an *in vivo* system, we discovered a cellular and molecular mechanism that maintains MT content and polarity in dendrites.

## Acknowledgments

We thank Lauren Cote and Jessica Feldman for providing a GFP11 expressing plasmid, an integrated gfp1-10 expressing *C. elegans* strain, and their scientific feedback. We thank members of the Shen lab and Feldman lab for their scientific feedback and discussion. We also thank Tim Stearns for his scientific feedback. **Funding:** K. Shen is an investigator in the Howard Hughes Medical Institute. This work was supported by NIH NINDS 1R01NS082208. K. Ori-McKenney is supported by R35 award (NIH NIGMS 5R35GM133688), and K. Eichel is supported by a HHMI Hanna H. Gray Fellowship.

## Author contributions

Conceptualization, X.L. and K.S.; Methodology, X.L., R.A., K.E., C.T., V.P., H.D., K.O.M. and K.S.; Investigation, X.L. and R.A.; Writing – Original Draft, X.L., and K.S.; Writing – Review & Editing, X.L., K.S., K.O.M., K.E., C.T., V.P. and H.D.; Funding Acquisition, K.S. and K.O.M.; Resources, X.L., K.E., K.S., R.A. and K.O.M.; Supervision, K.S.

## Declaration of interests

The authors declare no competing interests.

## Figure Legends

**Fig. S1 Microtubule organization and stability in PVD neurons.** A) Schematic of PVD morphologies in outgrowing and mature stage. B) Co-localization of GFP::TBA-1 and PTRN-1::tdTomato in the mature PVD primary dendrite. Red arrowheads, MT minus ends indicated by both GFP::TBA-1 intensity and PTRN-1::tdTomato localization. C) GFP::TBA-1 and UNC-116(G237A)::mCherry (rigor mutant) localization in both cell body and dendrites. D) Kymograph of GFP::TBA-1 and UNC-116(rigor)::mCherry dynamic in tertiary dendrites. E) Representative images of pcStar::TBA-1 localization before photo conversion, 0h and 1h after photo conversion. White box: region of photo conversion. F) Quantification of the normalized intensity of red channel in the photo converted region. n=14 animals. Scale bar, 5μm for distance, 10s for time.

**Fig. S2 Function of UNC-33 is conserved in different neurons.** A) Representative kymograph of EBP-2::GFP in distal and proximal regions of primary dendrites in both wild-type animals and *unc-33* mutants. B) Kymograph of GFP::TBA-1dynamics in UNC-33-AID tagged animals with or without auxin treatments. Blue lines, MT polymerization. C) Quantification of MT polarity in animals with different genotypes and treatments. P values are calculated by Brown-Forsythe and Welch ANOVA tests. D) GFP::TBA-1 localization in DA9 neuron in both wild-type animals and *unc-33* mutants. Orange lines, axons; blue lines, dendrites. Red arrowheads, MT gaps in both axons and dendrites. E) Quantification of gap number in DA9 neuron. F) Quantification of gap length in DA9 neuron. n number is the same as E). P values are calculated by Welch’s t test. Scale bar, 5μ m for distance, 10s for time.

**Fig. S3 MT dynamics in *unc-33* mutants.** A) Kymograph of GFP::TBA-1 dynamics in proximal dendrites of *unc-33* mutants. Blue arrows, MT sliding, red arrows, MT loss, yellow arrowhead, MT depolymerization. B) Kymograph of EBP-2::GFP in the posterior dendrite of wild-type animals and *unc-33* mutants. C) Quantification of MT polarity in posterior dendrite in wild-type animals and *unc-3*3 mutants. D-G) Quantification of MT dynamic speed in wild-type and *unc-33* mutants. H and I) Quantification of MT dynamicity in wild-type and *unc-33* mutants. J and K) Quantification of MT dynamic frequency in wild-type and *unc-33* mutants. L and M) Quantification of MT dynamic length in wild-type and *unc-33* mutants. N) Quantification of MT catastrophe frequency in wild-type and *unc-33* mutants. O) Quantification of MT sliding in animals with different genotypes. P values were calculated by Brown-Forsythe and Welch ANOVA tests. Scale bar, 5μm for distance, 10s for time. For Fig. S3C-3N, significance was analyzed by Welch’s t test.

**Fig. S4 Microtubule stability in *unc-33* mutants. A)** Representative images of pcStar::TBA-1 localization before photo conversion, 0h and 1h after photo conversion in *unc-33* mutants. White box: region of photo conversion. F) Quantification of the normalized ratio of red channel intensity between 1h after conversion and 0h after conversion in wild type animals and *unc-33* mutants in the photo converted region. n=14 for wild type and n=12 for *unc-33* mutants. C) Representative kymograph of TBA-1 and PTRN-1 dynamic in *unc-33* mutants. Scale bar, 5μm for distance, 10s for time.

**Fig. S5 UNC-33 forms patch-like structures in neurites.** A) Schematic of *unc-33* genomic structure and the insertion site of GFP or 3xGFP11. B) Endogenous UNC-33 expression pattern. Red dashed circle, PVD cell body. Scale bar, 5μm C) Endogenous UNC-33 localization in PVD and FLP axons labeled by combining UNC-33::GFP11 knock in worm and P*des-2*::GFP1-10 strain (left), and the intensity distribution of UNC-33 patches in axon (right). Red dashed boxes, axon region of PVD and FLP. Scale bar, 5μm for left panels, 2μm for the zoomed in images in right panels.

**Fig. S6 Mitochondria targeted UNC-33s sequesters free tubulins in PVD.** A) Co-localization of GFP::TBA-1 and Mito-mCherry::UNC-33s in PVD in wild type animals. Box 1, cell body region, box 2, dendritic region. Scale bar, 5μm. B) Co-localization of GFP::TBA-1 and Mito-mCherry::UNC-33s in PVD in *unc-33* mutants. Box 3, dendritic region and the kymograph of GFP::TBA-1 and mito-mCherry::UNC-33s dynamic. C) Kymograph of GFP::TBA-1 and Mito-mCherry::UNC-33s dynamic in the dendrite of *unc-33* mutants. Red arrows, MT rescue event occurs in the mito-mCherry::UNC-33s locus, blue box, MT sliding event. Scale bar, 5μm for distance, 10s for time.

**Fig. S7** UNC-33 patches are required for microtubule rescue. A) Colocalization of UNC-33::(int)GFP and UNC-44::FLPon-mScarlet in PVD, a1, UNC-33 and UNC-44 localization in the primary dendrite; a2, UNC-33 and UNC-44 localization in the high order branch. B) Second representative kymograph of UNC-33:(int)GFP and mCherry::TBA-1 dynamic in PVD tertiary dendrites in *unc-44(lf)* mutants. C) Second representative kymograph of UNC-33(E663K)::(int)GFP and mCherry::TBA-1 dynamic in PVD tertiary dendrites. Red arrows, MT shorting or loss. Scale bar, 5μm for distance, 10s for time.

## STAR Methods

### C. elegans strains

Worms were raised on NGM plates at 20°C using OP50 *Escherichia coli* as a food source and imaged at room temperature.

All the endogenous knock in worms were generated by CRISPR/Cas9 editing^66^. Transgenic strains were generated using standard microinjection techniques.

All the *C. elegans* strains used in this study are listed in table S1.

*C. elegans* staging and synchronization were performed as previously described^33^.

### Molecular biology

Plasmids, primers and gRNA sequence used to generate transgenic or knock in *C. elegans* strains in this study are listed in table S2. Plasmids were generated using T4 ligase or In-fusion cloning kit. gRNA used for CRISPR/Cas9 knock in were ordered from IDT. Specifically, the strain used for MT dynamic imaging is a transgenic strain with the expression of P*unc-86*::GFP::TBA-1a in *tba-1*(*ok1135*) mutant background to reduce the free tubulin background signal.

### Auxin Treatment

Before treatment, worms were cultured using standard method. Adult worms were collected for synchronization. Bleached worms were transferred onto unseeded plates and cultured at 20 degree overnight, then arrest L1s were transferred to OP50 seeded plates for around 25-26 hours at 20 degree, and worms were washed from the plates and transferred to 4mM auxin plates for 18 hours before time lapse imaging. Before transferred to auxin plates, some of the worms were examined by the microscope for the staging, worms were transferred to auxin plates when most of the worms finished primary growing or at the end of primary growing. Time cultured at 20 degree is slightly variable for worms with different genotype.

### Microscopy

Imaging of microtubule dynamic and protein localization was performed on an inverted Zeiss Axio Observer Z1 microscope equipped with a Yokogawa spinning disk, QuantEM:512SC Hamamatsu camera (set to 600 EM Gain), a Plan-Apochromat 100x/1.4 NA objective (Zeiss), 488 nm and 561 nm lasers, and controlled by MetaMorph Microscopy software (Molecular Devices). Imaging of microtubule array in wild-type and different mutants and UNC-33 localization were performed on spinning disk system (3i) with a CSU-W1 spinning disk (Yokogawa), 405-nm, 488-nm and 561-nm solid-state lasers, a C-Apochromat 63×/1.2 NA water-immersion objective, and a Prime95B camera (Photometrics).

Photo bleaching of UNC-33 signal and photo conversion of pcStar were performed on the spinning disk system (3i). Both bleaching and photo conversion were manipulated by 405-nm laser, bleaching was done by a 10% laser power while photo conversion was done with 0.1% laser power.

To image EBP-2 dynamic in outgrowing PVD neurites, arrest L1 stage wild-type animals were grown on OP50-seeded NGM plates at 20°C for 22-24 hours before imaging. *unc-33*, *unc-44* and *unc-119* mutants were grown on NGM plates at 20°C for 28-30 hours before imaging.

To get a better resolution for the MT dynamic in the growth cone region, GFP::TBA-1 animals were imaged 24-25 hours later when cultured on the OP50 plates at 20°C for wild type animals, in this stage primary dendrite is longer (passing the middle region of the worm) and MT number is reduced in the growth cone region. For the movies, images were taken every 300ms with a total number of 200 or 300 images.

To image the mature stage worms, L4 stage worms were selected to perform most of the imaging except for the UNC-33 localization worms. To get a better expression of *gfp*1-10 in the cell cytoplasmic, UNC-33::GFP11×3;ser-2P3::GFP1-10 worms were imaged at day1 adult stage. To visualize the UNC-33 localization specifically in PVD during development, a large region (around 30μm x 10 μm) before PVD growth cone was photo bleached, then UNC-33 localization and PVD morphology were imaged 10-20 minutes later.

### Image analysis and quantification

#### Images were processed and analyzed using ImageJ

Microtubule dynamic speed: Kymograph was made using the Reslice function in ImageJ, and the angle of the TBA-1 growing or shrinking line was measured as α, and then the speed was calculated by the formula: speed(μm/s)=0.109/0.3* cot α, for the time lapse imaging, time interval is 300ms between frames and scale for distance is 0.109μm/pixel.

For the line graph in Fig. 1J, the line distribution was made by Prism9(GraphPad) and the color was added using Adobe Illustrator manually.

Statistical calculations and graphing were done in Prism 9 (GraphPad). Co-localization analyze was done using the colocalization plug in in Fiji.

Any movie that showed a slight movement was aligned by ImageJ StackReg plugin before analysis.

### Microtubule assembly

Pig brains were obtained from a local abattoir and used approximately within four hours after death. Porcine brain tubulin was isolated using the high-molarity piperazine-*N*,*N*′-bis(2-ethanesulfonic acid) PIPES procedure and then labeled with biotin NHS ester or Dylight-405 NHS ester as described previously^67^. Taxol-lattice microtubules were polymerized for 30 min at 37 °C using 50 mM of unlabeled tubulin, 10 μM of biotin-labeled tubulin and 3.5 μM of Dylight-405-labeled tubulin in BRB80 (80 mM PIPES, 1 mM ethylene glycol-bis(2-aminoethylether)-*N*,*N*,*N*′,*N*′-tetraacetic acid (EGTA), 1 mM MgCl_2_, pH 6.9) supplemented with 2 mM GTP. The polymerized microtubules were then stabilized by addition of 20 μM taxol and incubated an additional thirty minutes. Microtubules were pelleted by centrifugation at 15,000 x g over a 25% sucrose cushion for ten minutes. The pellet was then resuspended in BRB80 supplemented with 10 μM taxol. GMPCPP seeds were made from a mixture of 647-tubulin, biotin-tubulin, and unlabeled tubulin that was diluted to a final tubulin concentration of 30 μM in BRB80 + 1 mM DTT. The mixture was then incubated with 1 mM GMPCPP (Jena Biosciences, NU-405) at 37 °C for 20 min, then pelleted through a 25% sucrose cushion for 10 min at 50,000 x g. The pellet was then resuspended in BRB80.

### Protein constructs and purification

For in vitro experiments, UNC-33 constructs were cloned into pET28a vectors using In-fusion. Constructs contain an N-terminal cassette consisting of a Strep-tag and a sfGFP fluorophore. Bacterial expression of these constructs was conducted in BL21 cells, grown at 37°C until an O.D. of 0.6 was reached. Cells were induced overnight at 18°C with 0.1mM IPTG. Cells were then pelleted and resuspended in lysis buffer (50 mM Tris pH 8, 150 mM K-acetate, 2 mM Mg-acetate, 1 mM EGTA, 10% glycerol) with protease inhibitor cocktail (Roche), 1 mM DTT, 1 mM PMSF, and DNAseI. Cells were subsequently dounced on ice and passed through an Emulsiflex press. 0.1% of Triton X-100 was added before clearing by centrifugation at *14,000 RPM* for 20 mins. Clarified lysate from bacterial expression was passed over a column with Streptactin XT Superflow resin (Qiagen/IBA). After incubation, the column was washed with five column volumes of lysis buffer and bound proteins were eluted with 50 mM D-biotin (CHEM-IMPEX) in lysis buffer. Eluted proteins were subsequently diluted in lysis buffer pH 8.5 and loaded on a 5mL HiTrap Q anion exchange column (GE Healthcare). Purified protein was eluted with a 0.1-0.4M NaCl gradient over 45 column volumes using a BioRad NGC Chromatography System. Peak fractions were collected, concentrated, and flash frozen in LN_2_.

### Single-molecule Mass Photometry

To prepare mass photometry chambers, microscope cover glasses (#1.5 24 × 50 mm, Deckgläser) were cleaned by a one hour sonication in Milli-Q H_2_O before a one hour sonication in isopropanol. Cover glasses were then coated with 0.1% Poly-L-Lysine before being briefly rinsed with Milli-Q H_2_O and dried by filtered air. CultureWell silicone gaskets (Grace Bio-Labs) were cut and washed in isopropanol and Milli-Q H_2_O before being dried by filtered air and placed onto cover glasses. Each cover glass and gasket setup provided four independent sample chambers. Samples and standard proteins were diluted in BRB80, freshly filtered with a 0.22 μm filter, and kept at room temperature. For calibration, standard proteins BSA (Sigma), Apoferritin (Sigma) and Thyoglobulin (Sigma) were diluted to 10-50 nM. Masses of these molecules were measured by OneMP instrument (Refeyn) and data was collected at an acquisition rate of 1 kHz for 100 s by AcquireMP (Refeyn). Data was subsequently analyzed by DiscoverMP (Refeyn).

### Total Internal Reflection Fluorescence Microscopy (TIRF-M)

TIRF microscopy experiments were performed using a 100X 1.49NA objective (Nikon) on a Nikon Ti-2 stand equipped with an Andor iXon EMCCD camera and four laser lines. The microscope was controlled by Nikon NIS Elements software. TIRF chambers were assembled from acid-washed coverslips and double-sided sticky tape as described (McKenney et al., 2014). Chambers were first incubated with 0.5 mg ml^−1^ PLL-PEG-biotin (Surface Solutions) for 5 minutes, followed by 0.5 mg ml^−1^ streptavidin for 5 minutes. Microtubules were diluted in BC Buffer (80 mM PIPES pH 6.8, 1 mM MgCl_2_, 1 mM EGTA, 1 mg ml^−1^ bovine serum albumin (BSA), 1 mg ml^−1^ casein). For taxol-stabilized microtubule experiments, BC buffer was supplemented with 10 μM taxol. Taxol or GMPCPP microtubules were flowed into the TIRF chamber post-streptavidin. To allow for adhesion, microtubules were incubated in the chamber for 5 minutes at room temperature and subsequently washed with TIRF assay buffer AB1 (60 mM HEPES pH 7.4, 50 mM potassium acetate, 2 mM MgCl_2_, 1 mM EGTA, 10% glycerol, 0.5% Pluronic F127, 0.1 mg ml^−1^ biotin-BSA, 0.2 mg ml^−1^ κ-casein) For taxol-stabilized microtubules, AB1 was supplemented with 10 μM taxol. For taxol-stabilized microtubule experiments, purified proteins were diluted to indicated concentrations in TIRF assay buffer and flowed into the TIRF chamber for imaging. For dynamic microtubule assays with GMPCPP seeds, sfGFP-UNC-33 was diluted to a final concentration of 50nM in assay buffer that also contained 15 uM of a mixture of 405-tubulin and unlabeled tubulin and 2 mM GTP. The entire mixture was flowed into the chamber containing GMPCPP seeds affixed to the coverslip then images were taken every 5 seconds for 5 minutes using an objective warmer set to 30 °C to capture growth and catastrophe of polymerizing microtubules. The experimental conditions were used in the absence of sfGFP-UNC-33. Resultant data was analyzed manually using ImageJ (Fiji).

## Supplemental information titles and legends

Table S1 *C. elegans* strains used in this study.

Table S2 Plasmids used in this study.

Table S3 gRNA and homology arm sequence used in this study.

## Notes

### Competing Interest Statement

The authors have declared no competing interest.

