## supplemental figures for "CRMP/UNC-33 maintains neuronal microtubule arrays by promoting individual microtubule rescue"

Fig. S1 Microtubule organization and stability in PVD neurons.

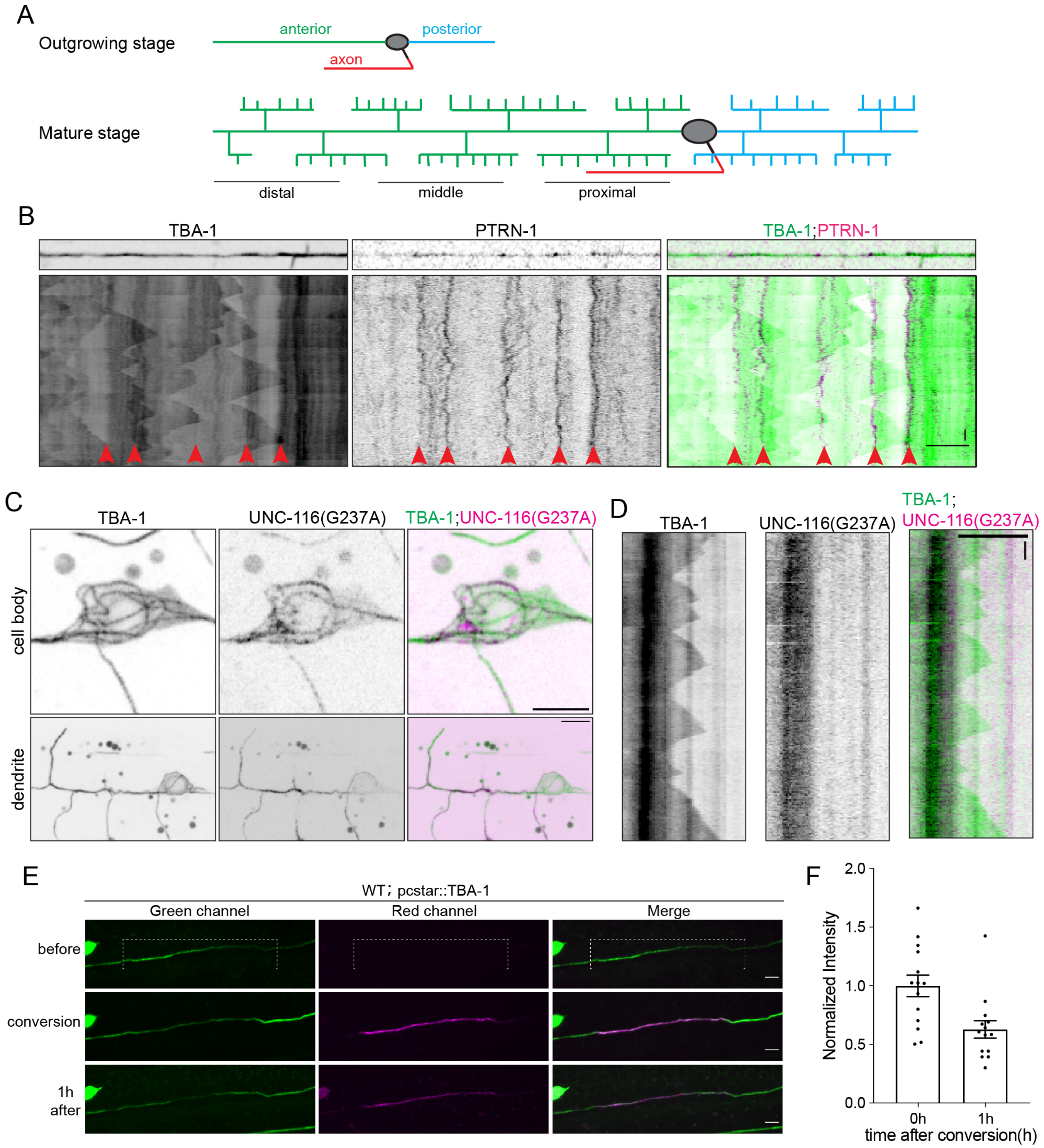

Fig. S2 Function of UNC-33 is conserved in different neurons.

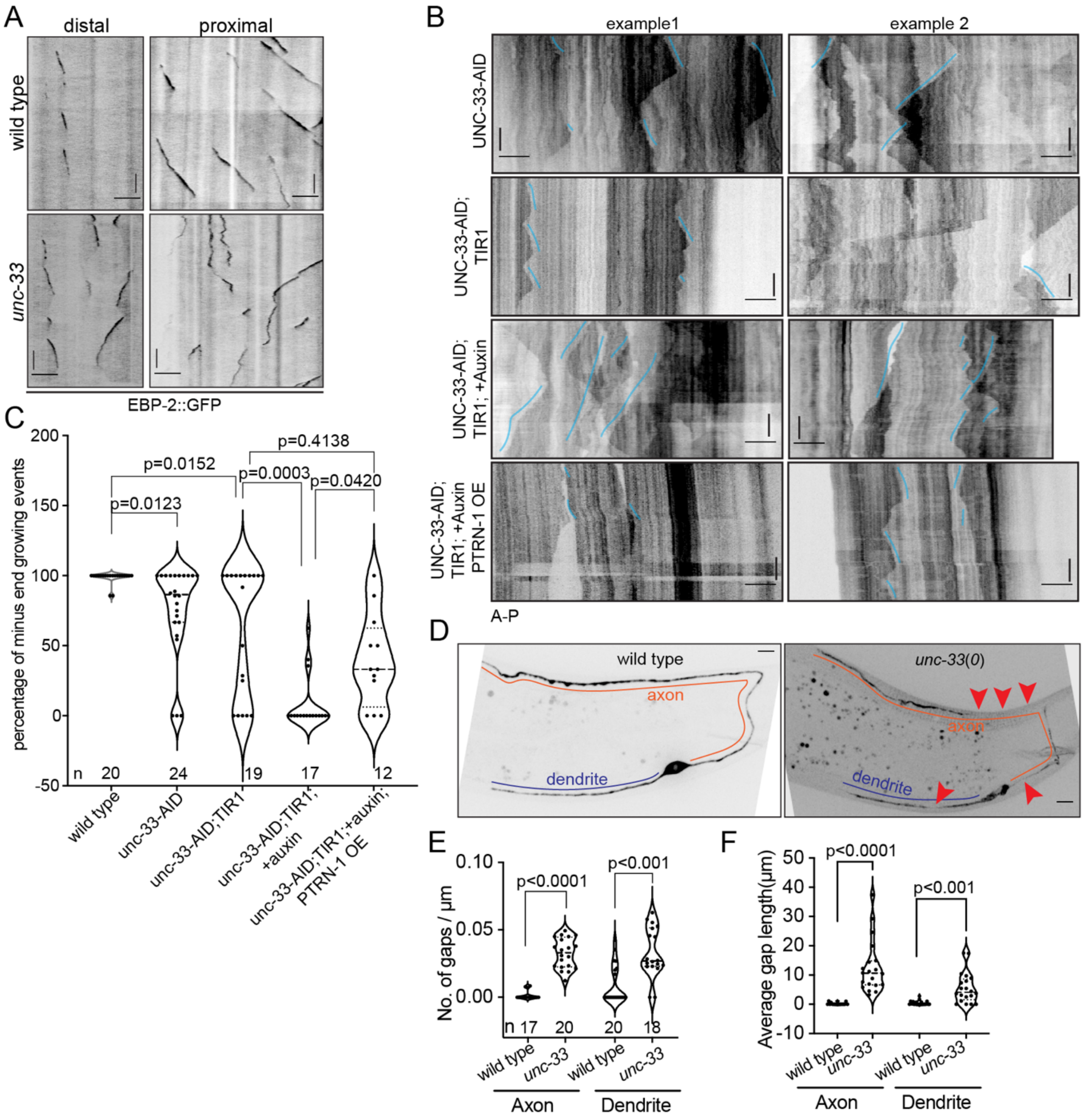

Fig. S3 MT dynamics in *unc-33* mutants.

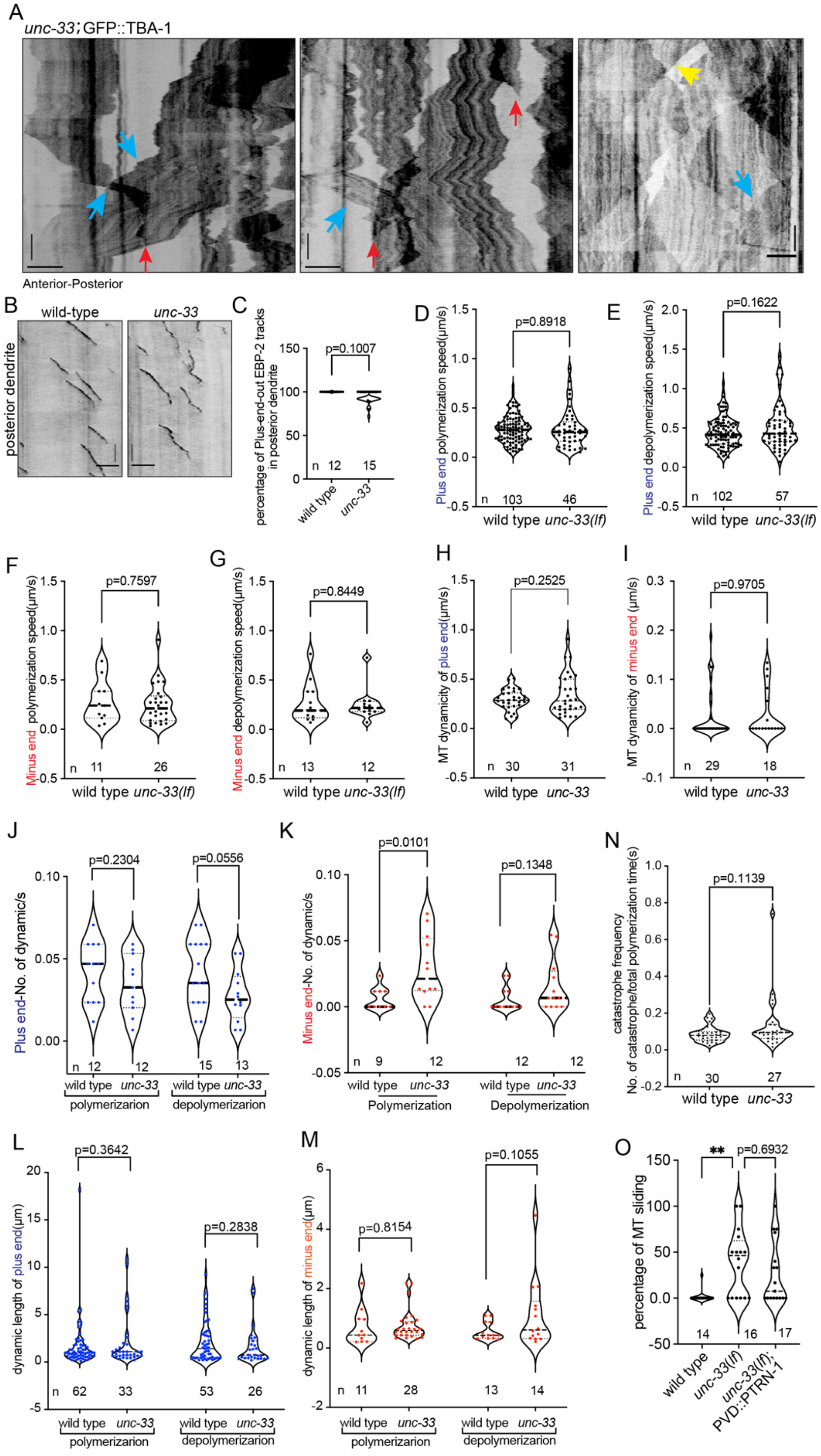

Fig. S4 Microtubule stability in *unc-33* mutants.

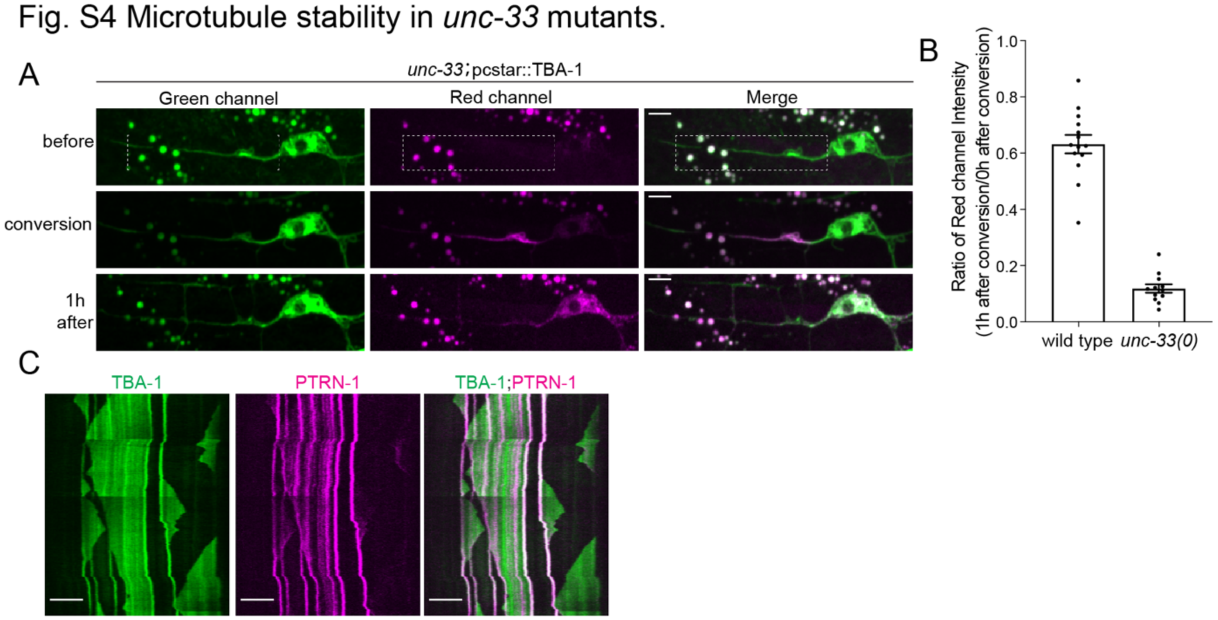


Fig. S5 UNC-33 forms patch-like structures in neurite.

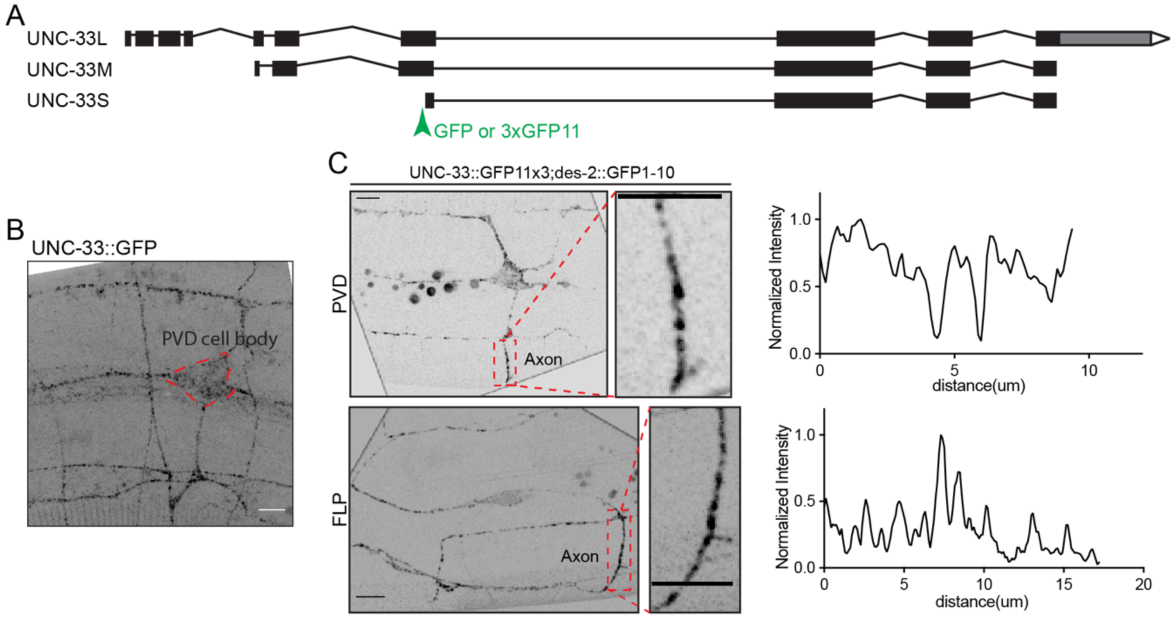

Fig. S6 Mitochondria targeted UNC-33S concentrates free tubulins in PVD.

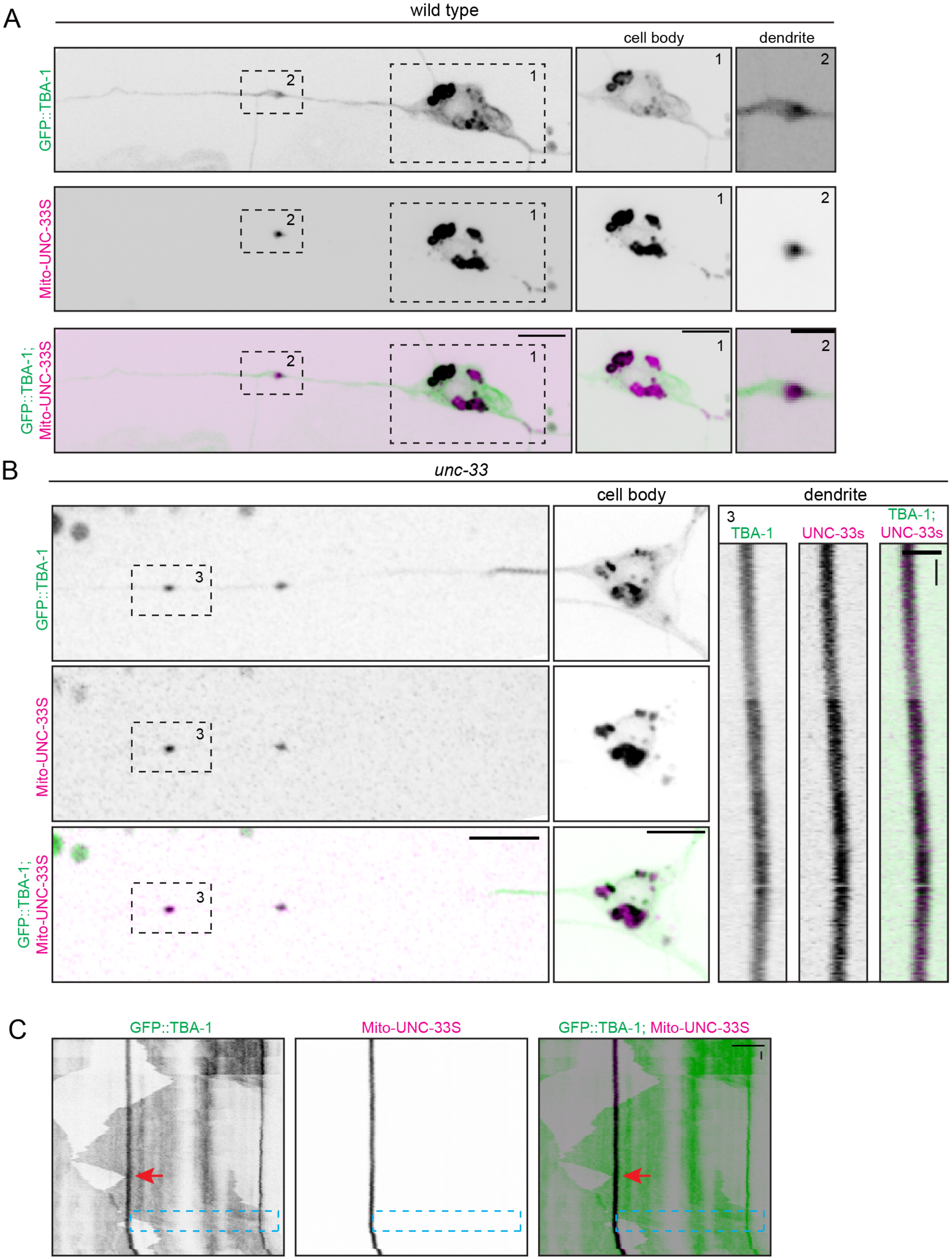

Fig. S7 UNC-33 patches are required for microtubule rescue.

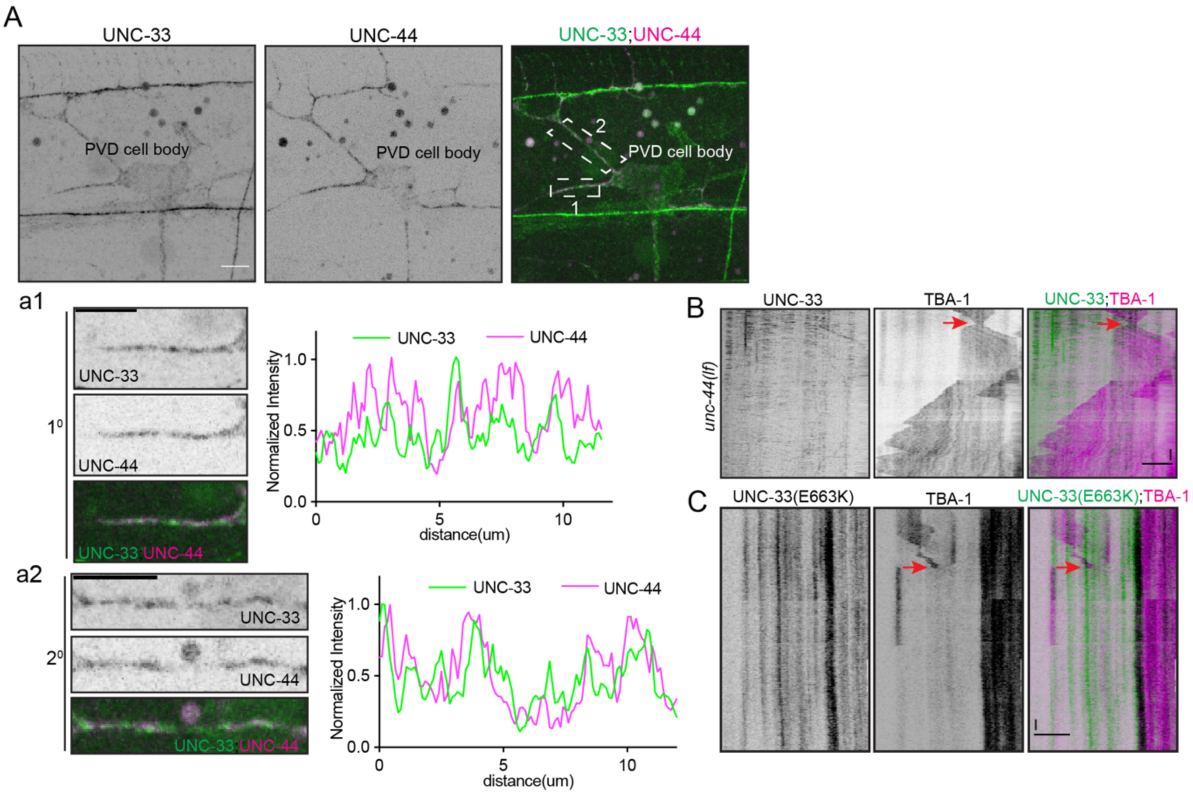
